# Hormetic nutrient stress promotes longevity by orchestrating histone acetylation on key lipid catabolism and antioxidant defense genes

**DOI:** 10.1101/2025.09.01.673564

**Authors:** Yifei Zhou, Fasih M. Ahsan, Sainan Li, Alexander A. Soukas

## Abstract

Exposure to low levels of environmental challenges, known as hormetic stress, such as nutrient deprivation and heat shock, fosters subsequent stress resistance and promotes healthy aging in later life. However, specific mechanisms governing transcriptional reprogramming upon hormetic nutrient stress remain elusive. In this study, we identified histone H3 lysine 27 acetylation (H3K27ac) as a crucial driver of transcriptomic adaptation to hormetic fasting. Beyond its immediate function of enhancing lipid catabolism for alternative energy sources, stress-induced H3K27ac activates lifelong antioxidant defenses, thereby reducing reactive oxygen species (ROS) produced by stress-induced fatty acid oxidation and their accumulation during aging. The increase in H3K27ac, mediated by pioneer factor PHA-4/FOXA and cooperating transcription factor NHR-49/HNF4, is crucial for lifespan extension under hermetic nutrient stress in *Caenorhabditis elegans*. Our findings establish H3K27ac as a key transcriptional switch that bridges nutrient status with transcriptomic reprogramming, underpinning the pro-longevity effects of hormetic fasting through orchestrating lipid catabolism and antioxidative defenses.

## Introduction

Environmental challenges, including temperature fluctuations, oxidative stress, and nutritional deprivation, pose significant risks to the survival and health of organisms. However, exposure to a low, sublethal dosage of stress known as hormetic stress, can promote organismal resistance against subsequent stress exposure and longevity by activating defense responses^1^. Caloric restriction (CR), likely the most experimentally and clinically tractable way of inducing hormesis, has been demonstrated to extend lifespan across various species and is correlated with enhanced healthspan in humans^2–4^. Different CR regimens, such as intermittent fasting, reducing food intake, and deprivation of specific nutrients, have been demonstrated to extend lifespan and health span via diverse mechanisms across species^5^. In *C. elegans*, where many CR regimens have been established to positively influence healthy aging, we previously demonstrated that a 24-hour period of food deprivation during early adulthood induces hormesis and promotes longevity, in a manner dependent on an AMP-dependent protein kinase (AMPK) / nucleoporin 50 (NUP50, NPP-16 in *C. elegans*) signaling axis^6^. Yet, the molecular mechanism of how NPP-16/NUP50 regulates transcription and chromatin plasticity remains unclear. Demonstrating the connection between energy sensing and chromatin accessibility could also reveal vital modulators governing the adaptive transcriptional response to hormetic nutrient stress that promotes healthy aging and alleviates aging-related disorders.

Because a hyperactivated stress response can be detrimental to organismal health and survival^7^, the dosage of hormetic stress must be precisely controlled to confer overall benefits for healthy aging. Upon nutrient and energetic deprivation, catabolic pathways are activated and shift from glucose to fat consumption to maintain energy production and survival^8^. As a result of activated fatty acid oxidation (FAO), the dominant approach breaking down fatty acids into acetyl-CoA, mitochondrial reactive oxygen species (ROS) generation is consequently accelerated due to the activated electron transport chain (ETC), thereby leading to cell death^9^. As a group of metabolites critical for intracellular redox homeostasis and various signaling pathways, ROS play crucial roles in signal transduction and fundamental cellular processes; however, excessive ROS is harmful to DNA, proteins, and organelles^10^. Its homeostasis is thereby critical for healthy aging and longevity^11^. As an adaptive response and a negative feedback, oxidative stress caused by activated lipid catabolism upon nutrient deprivation is mitigated at least partially by post-translational activation of superoxide dismutase (SOD2) in mice^12^. Nevertheless, the comprehensive spectrum of transcriptional reprogramming that orchestrates FAO and antioxidant defense in response to nutrient stress remains incompletely understood.

Epigenetic modification serves as a critical regulatory element for transcription and has been shown to be dysregulated with age and to play a vital role in aging-related disorders^13, 14^. Growing evidence suggests that epigenetic modifications are altered in response to various stresses and are essential for transcriptional adaptation in stress-associated longevity paradigms^15, 16^. Depletion of H3 lysine 27 (H3K27) methylation and subsequent enhanced acetylation (H3K27ac), well-established features of enhancers that positively impact transcriptional activity, are necessary for the mitochondrial unfolded protein response (UPR^mito^) when the ETC is inhibited^15^. H3K27 demethylation plays a crucial role in the heat shock response, facilitating the transcriptional activation of HSP70 chaperones^16^. H3K27ac also transduces effects of transcription factors CREB1 and PPAR for the transcriptional adaptation in response to starvation in mouse liver^17^. However, the mechanism by which the genomic site-specificity of epigenetic modifications responds to diverse stressors to drive distinct outcomes remains elusive. As a hallmark of aging, dysregulated epigenetic modifications are closely associated with and causally linked to the aging process^18^. Thus, long-term effects of epigenetic rewriting caused by acute hormetic stress are of great interest and potential therapeutic relevance.

In this study, we identify H3K27ac as the transcriptional switch that connects energetic availability and chromatin accessibility, facilitating metabolic rewiring and antioxidant defense for longevity. JMJD-3.1/JMJD3 and CBP-1/CBP, two vital H3K27ac modulators, are required for the lifespan extension prompted by elevating NPP-16/NUP50 expression levels, a longevity paradigm which molecularly mimics energetic stress^6^. The parallel reduction in H3K27 methylation and increase in H3K27ac modulated by these two key enzymes is also necessary for the pro-longevity effect of hormetic fasting in early life. Transcriptomic analysis of responses to hormetic nutrient stresses occurring early in life revealed that H3K27ac not only acutely and temporarily activates lipid catabolism but also activates persistent, lifelong antioxidant defenses, therefore improving resistance against stress and extending lifespan in *C. elegans*. The site-specificity of H3K27ac modulation across the genome is determined by pioneer factor PHA-4/FOXA and cooperating transcription factor NHR-49/HNF4. These findings reveal a mechanism that precisely coordinates metabolic rewiring and antioxidant defenses to promote healthy aging in response to hormetic nutrient stress and activation of the AMPK-NUP50-H3K27ac signaling axis.

## Results

### JMJD-3.1 and CBP-1 are required for longevity promoted by NPP-16 activation

To identify factors controlling the metabolic rewiring and pro-longevity effects of hormetic fasting, we performed a reverse genetic screen targeting epigenetic modulators using RNA interference (RNAi). Given that 24-hours of food deprivation could affect the efficiency of RNAi, which is conducted by feeding worms with *E. coli* expressing dsRNA against the genes of interest^19, 20^, we performed the epigenetic RNAi screen on a transgenic strain overexpressing NPP-16/NUP50 (*npp-16OE*), which phenocopies the transcriptomic alteration and lifespan extension upon hormetic nutrient stress^6^. We screened for epigenetic factor RNAi which blunts the survival rate of *npp-16OE* worms at day 20 adulthood (D20) when the wild-type (WT) worms are mostly dead, using *npp-16OE* as a proxy for the effects of hormetic fasting (Fig. 1a). We further tested any RNAi hits which shorten lifespan in *npp-16OE* worms for their impact on *npp-16OE-*mediated activation of the transcriptional reporter of *acs-2/*ACSF2, a widely used reporter of longevity associated with energetic stress^6, 21–25^. As epigenetic modulators could be necessary for survival and development of *C. elegans*^26, 27^, their RNAi might shorten the lifespan of WT worms in parallel to the effects of *npp-16OE*. Consequently, and in order to examine for true epistatic factors, we examined the full survival curve of WT and *npp-16OE* worms for RNAi candidates that met the two aforementioned screening criteria. To avoid potential developmental defects, worms were fed HT115 bacteria carrying the empty RNAi vector (EV) until the fourth larval stage (L4), then switched to the indicated clones for lifespan assessment following post-developmental RNAi knockdown (Fig. 1a).

**Fig. 1.**
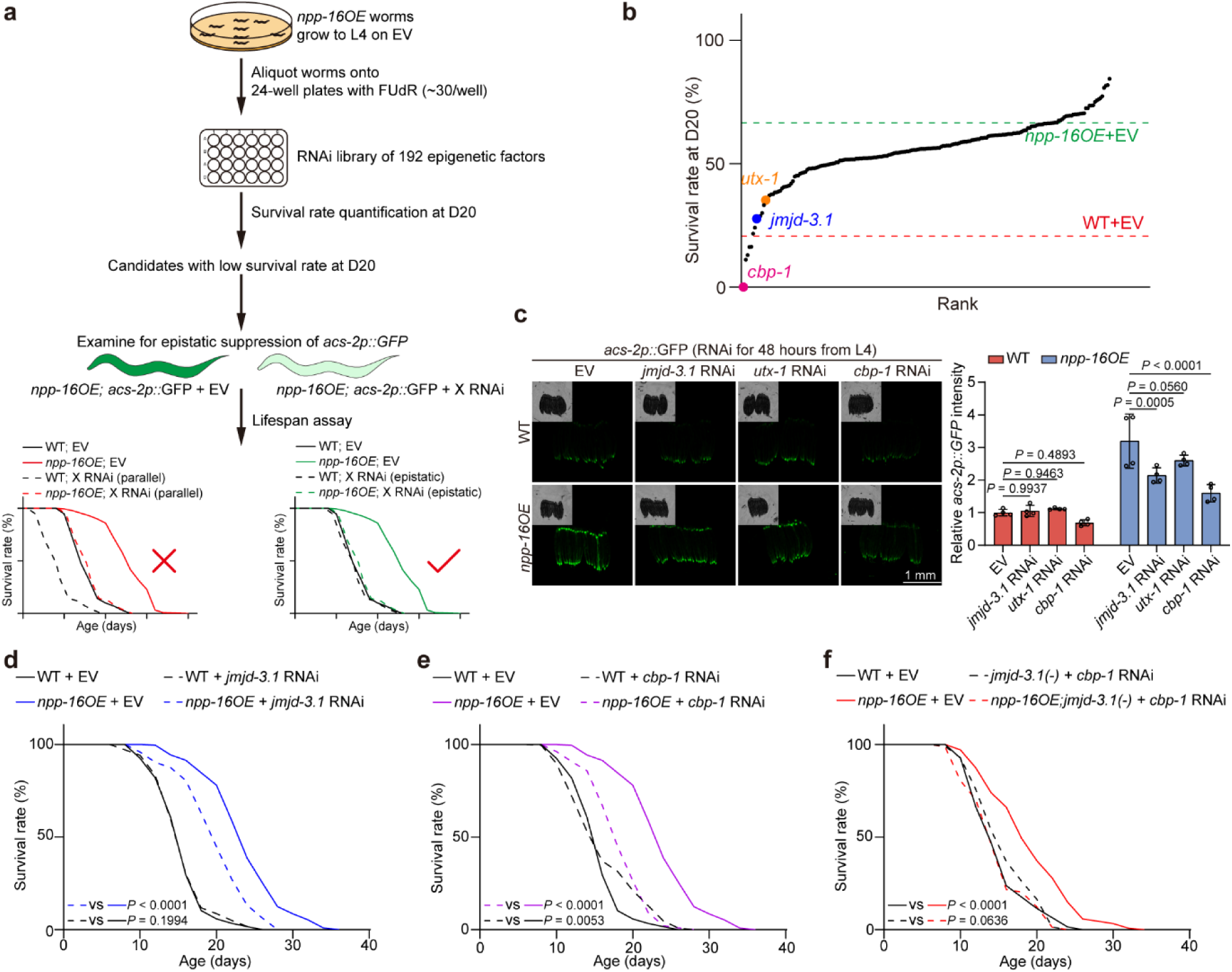
JMJD-3.1 and CBP-1 are required for the lifespan extension by NPP-16 activation. a, A depiction showing the RNAi screen strategy in the worms overexpressing NPP-16 (*npp-16OE*) for the epigenetic factors necessary for the activated lipid catabolism and extended lifespan. FUdR: 5-fluorodeoxyuridine. X RNAi indicates a potential downstream hit from the screen. **b,** The survival rate of *npp-16OE* worms fed with HT115 bacteria from the epigenetic factors RNAi library at day 20 of adulthood (D20). The D20 survival rates of WT and *npp-16OE* worms treated with the empty vector (EV) are indicated by red and green dashed lines as negative and positive controls, respectively. The H3K27ac regulators are highlighted. See Supplementary Data 1 for more information. n=2 independent experiments. **c,** Post-developmental RNAi of *jmjd-3.1* and *cbp-1* inhibits the activated *acs-2p::GFP* reporter in *npp-16OE* worms. L4, the fourth larval stage. Scale bar: 1 mm. n=4 independent experiments. **d,e**, RNAi of *jmjd-3.1* (**e**) and *cbp-1* (**f**) suppresses the lifespan extension significantly in *npp-16OE* worms. Worms were treated with RNAi from L1. **f**, Inhibited H3K27ac caused by *jmjd-3.1(-)* mutation and *cbp-1* RNAi blunts the lifespan extension in *npp-16OE* worms. See also Supplementary Data 3 for independent biological replicates and summary lifespan statistics. Error bars represent SD. Statistical significance was calculated by two-way ANOVA (**c**) and and Mantel-Cox Log Rank (**d-f**).

By quantifying the survival rate at D20, when the survival rates are 66.6% and 20.7% in WT and *npp-16OE* worms on empty vector (EV) plates, respectively (Fig. 1b), we identified a group of genes required for the longevity phenotype in *npp-16OE* worms (Fig. 1b and Supplementary Data 1), including *jmjd-3.1*/JMJD3 and *utx-1*/UTX, two H3K27 demethylases^28, 29^, and *cbp-1*/CBP, the ortholog of the dominant functional lysine acetyltransferase CBP/p300 in *C. elegans*^30^ (Extended Data Fig. 1a). H3K27 di-and tri-methylation is a transcriptional repressor, and H3K27 demethylation is required for subsequent H3K27ac for transactivation at promoters and enhancers^31, 32^. Concordantly, L4-onset RNAi knockdown of *jmjd-3.1* and *cbp-1* significantly suppresses the induced *acs-2p::GFP* reporter in *npp-16OE* worms, whereas *utx-1* RNAi has a milder and non-significant effect (Fig. 1c). Moreover, L1-onset RNAi knockdown of *jmjd-3.1* and *cbp-1* also partially suppresses the lifespan extension of *npp-16OE* individually without appreciable shortening of wild type lifespan (Fig. 1d,e). Consistent with *jmjd-3.1* RNAi knockdown, *jmjd-3.1(-)* mutation also significantly attenuates the longevity phenotype of *npp-16OE* worms (Extended Data Fig. 1c). We then depleted H3K27ac by treating a loss-of-function mutant of *jmjd-3.1* (*jmjd-3.1(-)*) with *cbp-1* RNAi that inhibits two genes together. As we hypothesized based upon the results of our screen, activating NPP-16 fails to extend the lifespan in worms with impaired H3K27ac (Fig. 1f). In aggregate, H3K27ac modulators are necessary for activation of *acs-2* expression and concomitant longevity associated with increased expression of NPP-16/NUP50.

Since there are four H3K27 demethylases orthologous to JMJD3 and UTX/UTY in *C. elegans* (Extended Data Fig. 1a), we examined the effects of these other demethylases on *acs-2p::GFP* reporter expression and lifespan in *npp-16OE* worms*. jmjd-3.2* and *jmjd-3.3* RNAi cause a milder decrease of *acs-2p::GFP* reporter expression when compared to *jmjd-3.1* RNAi (Extended Data Fig. 1b). Strongly in contrast to *jmjd-3.1*, RNAi of *jmjd-3.2* and *jmjd-3.3* do not influence the lifespan extension of *npp-16OE* animals (Extended Data Fig. 1d,e). Unlike other H3K27 demethylases, *utx-1* RNAi leads to a significant lifespan extension in WT worms^33^, and does not significantly impact the pro-longevity impact of *npp-16OE* (Extended Data Fig. 1f), implying a distinct and parallel role of UTX-1/UTX in aging. These results indicate that JMJD-3.1 is the dominant H3K27 demethylase necessary for the metabolic and longevity-promoting effects of *npp-16OE* and further suggest an essential role in the hormetic response to nutrient stress.

### NPP-16 overexpression elevates H3K27ac for the transactivation of lipid catabolism

As a stress response to food deprivation, NPP-16/NUP50 levels increase, leading to transcriptional induction of *acs-2*/ACSF2, *lipl-1*/LIPA, *lipl-2*/LIPA, and *lipl-3*/LIPA and subsequent activation of lipid catabolism^6, 21, 34^. This transcriptional activation is blocked when H3K27ac is inhibited by *jmjd-3.1(-)* mutation and *cbp-1* RNAi knockdown (H3K27ac(-), Fig. 2a). The reduced fat mass prompted by NPP-16 activation is also blunted in worms with impaired H3K27ac (Fig. 2b and Extended Data Fig. 2a), further confirming that JMJD-3.1/JMJD3 and CBP-1/CBP are required to activate lipid catabolism in *npp-16OE* worms. However, neither *npp-16* null mutation (*npp-16(-)*) nor *npp-16OE* has a significant effect on global H3K27ac levels in worms at day 1 of adulthood (Extended Data Fig. 2b). We therefore suspected that H3K27ac might be only enhanced on the promoters of specific genes for metabolic rewiring without global alteration. By chromatin immunoprecipitation (ChIP), we find that H3K27ac is indeed increased by *npp-16OE* on the promoter of *acs-2* (Fig. 2c), whereas it is unchanged at the 3’-UTR of *acs-2* and the *act-1* promoter (Fig. 2c and Extended Data Fig. 2c). These data suggest that factors that modulate H3K27ac may be the effector arm by which increased NPP-16/NUP50 promotes transactivation of lipid catabolism and longevity.

**Fig. 2.**
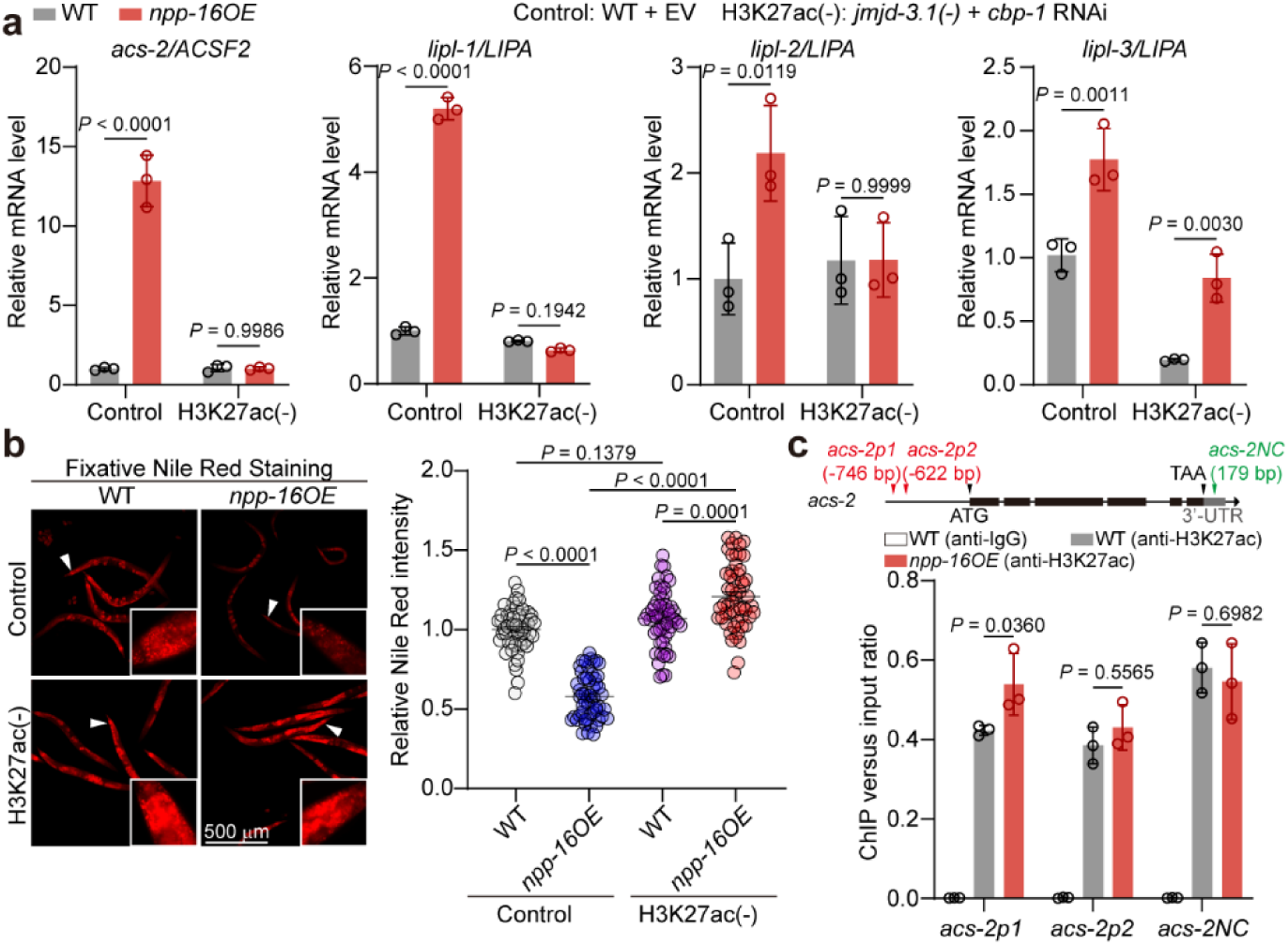
NPP-16/NUP50 transactivates the lipid catabolic genes through elevating H3K27ac on promoters. **a**, qRT-PCR analyses reveal that the induced transcription of lipid catabolic genes *acs-2*, *lipl-1*, *lipl-2*, and *lipl-3* is rescued by H3K27ac inhibition (H3K27ac(-)) conducted by the combination of *jmjd-3.1(-)* knockout and *cbp-1* RNAi knockdown. n=3 independent experiments. **b**, Neutral lipid storage indicated by fixative Nile red staining reveals that the reduced fat mass in *npp-16OE* worms is blocked by H3K27ac(-) at day 1 adulthood. Scale bar: 500 μm. n=3 independent experiments. **c**, ChIP-PCR analyses reveal that *npp-16OE* is sufficient to elevate H3K27ac on the promoter regions of *acs-2*. The flanking sequences at or near the 3’-UTR serve as the negative controls (*acs-2NC*) for ChIP-qPCR. n=3 independent experiments. Error bars represent SD (**a**,**c**) and SEM (**b**). Statistical significance was calculated by two-way ANOVA (**a**,**c**) and one-way ANOVA (**b**).

### Hormetic fasting promotes transcriptomic reprogramming of lipid catabolism and antioxidant defenses in a H3K27ac-dependent manner

We were next interested in the role of H3K27ac modulators in longevity caused by hormetic nutrient stress. However, single RNAi knockdown of *jmjd-3.1* and *cbp-1* only has mild impacts on the lifespan extension achieved through hormetic fasting for 24 hours at day 1 of adulthood (Extended Data Fig. 3a,b), which may be due to the inefficiency of RNAi during food depletion^19, 20^. *jmjd-3.1(-)* mutation also only partially suppresses the lifespan extension of hormetic fasting (Extended Data Fig. 3c), showing that inhibiting H3K27 demethylation alone cannot block the longevity phenotype of hormetic fasting. We next tested the pro-longevity effects of hormetic fasting in the worms with defective H3K27ac by treating *jmjd-3.1(-)* mutants with *cbp-1* knockdown (Fig. 3a). In keeping with our hypothesis, upon fasting at day 1 of adulthood, H3K27ac(-) worms exhibited a comparable lifespan with those fed ad libitum (AL, Fig. 3a,b), indicating that H3K27ac is necessary for the lifespan extension of hormetic fasting.

**Fig. 3.**
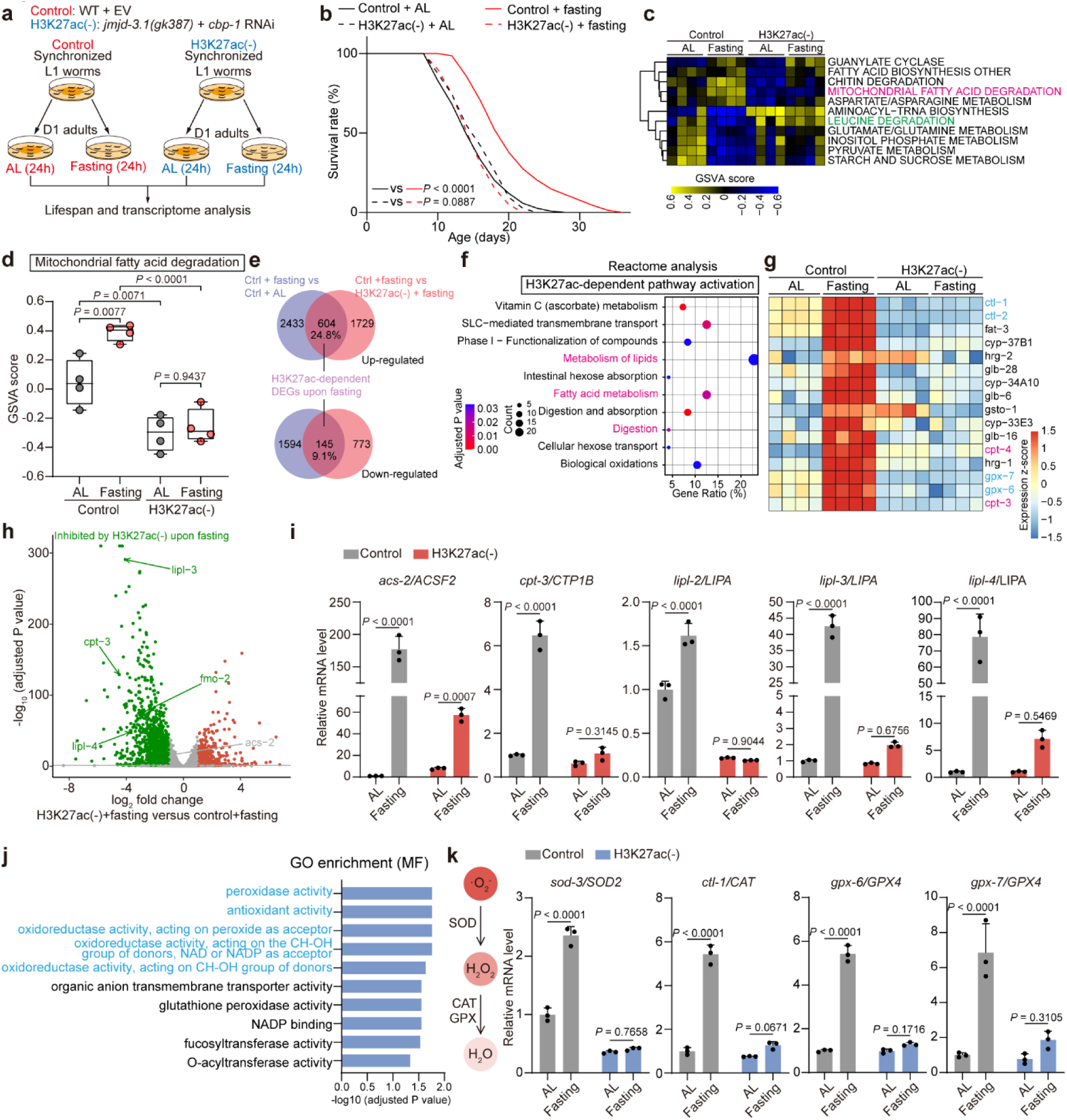
JMJD-3.1 and CBP-1 are necessary for the transcriptomic alteration and longevity promotion of hormetic fasting. **a**, A depiction showing the H3K27ac inhibition (H3K27ac(-)) and feeding strategy for the following lifespan analyses and RNA sequencing (RNA-seq). H3K27ac inhibition is conducted by the combination of *jmjd-3.1(gk387)* knockout and *cbp-1* RNAi knockdown. AL, ad libitum feeding. **b**, H3K27ac inhibition blunts the lifespan extension of hormetic fasting for 24 hours at D1. See also Supplementary Data 3 for independent biological replicates and summary lifespan statistics. **c-d**, GSVA analysis of the RNA-seq data reveals that mitochondrial fatty acid degradation is significantly activated by fasting and is blocked by H3K27ac(-). **e**, A Venn diagram showing the number of differentially expressed genes (DEGs) in control (Ctrl) worms subjected to fasting versus AL, and in fasted control worms versus fasted H3K27(-) worms. The overlapping DEGs from these two sets represent the H3K27ac-dependent transcriptomic alteration upon fasting. **f**, Reactome analysis of RNA-seq data shows that pathways relevant to lipid catabolism are activated by fasting in a H3K27ac-dependent manner. **g**, A heat map showing the representative DEGs induced by fasting depending on H3K27ac. Lipid catabolic and antioxidant genes are highlighted in magenta and blue, respectively. **h**, A volcano plot highlights the DEGs from RNA-seq analyses of H3K27ac(-) worms versus control worms upon fasting. Red and green dots represent the significantly activated and inhibited DEGs by H3K27ac(-) in response to hormetic fasting. **i**, qRT-PCR analyses reveal that lipid catabolic genes participating in lysosomal lipolysis and fatty acid oxidation are induced by fasting in a H3K27ac-dependent manner. n=3 independent experiments. **j**, GO-term of molecular function (MF) overrepresentation analysis of the 604 H3K27ac-dependent up-regulated DEGs by fasting. Antioxidation-relevant functions are highlighted in blue. **k**, qRT-PCR analyses reveal that key enzymes participating in ROS degradation are elevated by fasting in a H3K27ac-dependent manner. n=3 independent experiments. See Supplementary Data 2 for detailed statistics from RNA-seq (**c**, **f-h**, and **j**). Error bars represent SD. Statistical significance was calculated by Mantel-Cox Log Rank (**b**), empirical Bayes testing with Benjamini-Hochberg correction for multiple comparisons using limma()(**d**), and two-way ANOVA (**i**,**k**).

To further investigate transcriptomic alteration of H3K27ac in response to nutrient stress, we performed bulk RNA sequencing (RNA-seq) with the RNA extracted from control and H3K27ac(-) worms subjected to different feeding strategies (Fig. 3a). By performing gene set variation analysis (GSVA)^35^, we found that mitochondrial fatty acid degradation is significantly activated by fasting in control worms but blunted by H3K27ac inhibition (Fig. 3c,d and Extended Data Fig. 2d), which is similar to transcriptional alterations previously reported in response to short-term starvation and activation of NPP-16^6, 21^. Genes encoding leucine catabolic enzymes are mildly inhibited by fasting, and this shift is attenuated by H3K27ac depletion (Extended Data Fig. 3d,e). By defining differentially expressed genes (DEGs) as those with a log_2_ fold change greater than 1.0 and an adjusted *P* value less than 0.05, we identified 604 and 145 genes that are significantly increased and decreased respectively by fasting in a H3K27ac-dependent manner (Fig. 3e and Supplementary Data 2). Since H3K27ac is a marker for enhancer and promoter transcriptional activation^36^, we then principally focused on the 604 up-regulated DEGs that is about 24.8% of the up-regulated transcriptome upon food deprivation (Fig. 3e). As expected, we found that the pathways relevant to fatty acid metabolism were activated by fasting and prevented by H3K27ac(-) through Reactome pathway overrepresentation analysis and annotation (Fig. 3f)^37, 38^. The top H3K27ac-depedent DEGs upon hormetic fasting contain many lipid catabolic genes involved in lipolysis and fatty acid oxidation (Fig. 3g,h and Extended Data Fig. 4b), which was confirmed by qRT-PCR, such as the aforementioned nematode fatty acid CoA synthetase *acs-2/ACSF2*, *cpt-3/CPT1B* which encodes carnitine palmitoyl transferase in *C. elegans*, *lipl-2/LIPA*, *lipl-3/LIPA*, and *lipl-4/LIPA*, the orthologues of lysosomal acid lipases (Fig. 3i). We next compared transcriptional changes associated with hormetic starvation and metabolic rewiring and longevity promotion in *npp-16OE* worms (Fig. 1) by overlapping the transcriptomes of fasting and *npp-16OE*^6^. Suggestive of common mechanistic underpinnings, 35.8% of H3K27ac-dependent DEGs are also under the control of activated NPP-16 (Extended Data Fig. 4a).

Interestingly, besides lipid catabolism, we also found that many enzymes in ROS scavenging and buffering are induced by fasting and blunted by H3K27ac impairment (Fig. 3g). By GO-term analysis^39, 40^, we also identified i) peroxidase activity, ii) antioxidant activity, iii) oxidoreductase activity, and iv) longevity-regulating pathway as terms promoted by fasting in an H3K27ac-dependent manner (Fig. 3j and Extended Data Fig. 3f-h). We then hypothesized that the enhanced antioxidant defense could eliminate the excess and maladaptive ROS produced by enhanced fatty acid oxidation. During ROS scavenging, the dismutation of superoxide anion is catalyzed into less reactive hydrogen peroxide by superoxide dismutase (SOD). Hydrogen peroxide is then broken down into harmless water by catalase (CAT) and glutathione peroxidase (GPX, Fig. 3k)^41^. Strikingly, *sod-3/SOD2*, *clt-1/CAT*, *ctl-2/CAT*, *gpx-6/GPX4*, and *gpx-7/GPX4* are all increased by fasting in a H3K27ac-dependent manner (Fig. 3g,k). Consistent with these findings, antioxidant mRNAs corresponding to genes *gpx-6* and *gpx-7* are also activated in *npp-16OE* worms (Extended Data Fig. 3i). In aggregate, hormetic fasting activates the transcription of antioxidant responses in an H3K27ac-dependent manner in parallel to lipid catabolism. We hypothesize that antioxidant defenses may attenuate ROS accumulation prompted by fatty acid oxidation activated during fasting.

### Hormetic fasting specifically activates H3K27ac-dependent lipid and antioxidant stress defenses

Given that JMJD-3.1/JMJD3 and CBP-1/CBP are also responsible for the adaptive UPR^mito^ activation upon ETC inhibition^15^ and JMJD-3.1/JMJD3 is required for the heat shock response^16^, we next wondered about the transcriptomic overlap between nutrient stress and other stress responses, and whether H3K27ac can be specifically deployed to response genes dependent upon the specific stress encountered. Indeed, only 11.6% and 3.0% of H3K27ac-dependent DEGs upon fasting overlap with the up-regulated transcriptome of *cco-1* RNAi and *cbp-1*-dependent DEGs upon ETC inhibition, respectivetly^15^ (Extended Data Fig. 4c,d). Moreover, the genes markedly induced during heat shock response and UPR^mito^ are largely unaffected in response to fasting (Supplementary Data 2). Autophagy is a well-known adaptation to energetic and nutrient stress, activated to maintain proteostasis^42^. Although some critical autophagic genes are induced by fasting, such as *lgg-1/LC3*, *pha-4/FOXA*, and *sqst-1/SQSTM*, they are not dependent upon H3K27ac (Extended Data Fig. 4e), suggesting a dispensable role of H3K27ac in autophagy activation in response to nutrient challenge.

Concordantly, among the fluorescent reporters of various stress responses, *acs-2p::GFP* reporter is the only one significantly activated by hormetic fasting (Extended Data Fig. 4f). In contrast, other reporters *hsp-16.2p::GFP*, *hsp-6p::GFP*, and *hsp-4p::GFP*, representing the activity of UPR^mito^, endoplasmic reticulum unfolded protein response (UPR^ER^), and heat shock response respectively, are unchanged upon fasting (Extended Data Fig. 4g-i). These findings indicate an unappreciated H3K27ac-dependent transcriptomic alteration controlled by hormetic fasting that is distinct from other stress responses.

### Hormetic fasting promotes H3K27ac at lipid catabolic and antioxidative genes

The global H3K27ac level is elevated upon ETC inhibition in a JMJD-3.1- and CBP-1-dependent manner^15^. However, we do not observe a similar increase in fasted worms (Extended Data Fig. 4j), suggesting that the H3K27ac landscape might be redistributed in a site-specific manner to activate expression of fasting response genes without a global alteration of H3K27ac. This finding is concordant with the unchanged H3K27ac in *npp-16OE* worms (Extended Data Fig. 2b). To verify the action of JMJD-3.1 and CBP-1 on lipid catabolic genes specifically, we detected the H3K27ac level at the promoter regions of lipid catabolic genes upon fasting. In line with the mRNA induction, H3K27ac levels are elevated by hormetic fasting on the promoter regions of *lipl-3* and *acs-2.* In contrast, it is unchanged on their 3’-UTRs and *act-1* promoter (Fig. 4a,b and Extended Data Fig. 4k). As predicted, the fasting-induced H3K27ac on *lipl-3* and *acs-2* promoter is ablated in worms with inhibited JMJD-3.1 and CBP-1 (Fig. 4a,b), indicating they are the dominant modulators of the H3K27ac induction on lipid catabolic genes in fasted animals. We also find a significant increase of H3K27ac on the promoters of antioxidant genes *ctl-1/2*, *sod-3*, and *gpx-7*, which is blocked by *jmjd-3.1(-)* mutation and *cbp-1* RNAi knockdown (Fig. 4c,d, and Extended Data Fig. 4l). We also observed that H3K27ac increases at the 3’-UTR of *sod-3* and *gpx-7* in a manner dependent on JMJD-3.1 and CBP-1. In aggregate, these findings show that hormetic fasting induces H3K27ac on lipid catabolism and antioxidant genes in a JMJD-3.1- and CBP-1-dependent manner.

**Fig. 4.**
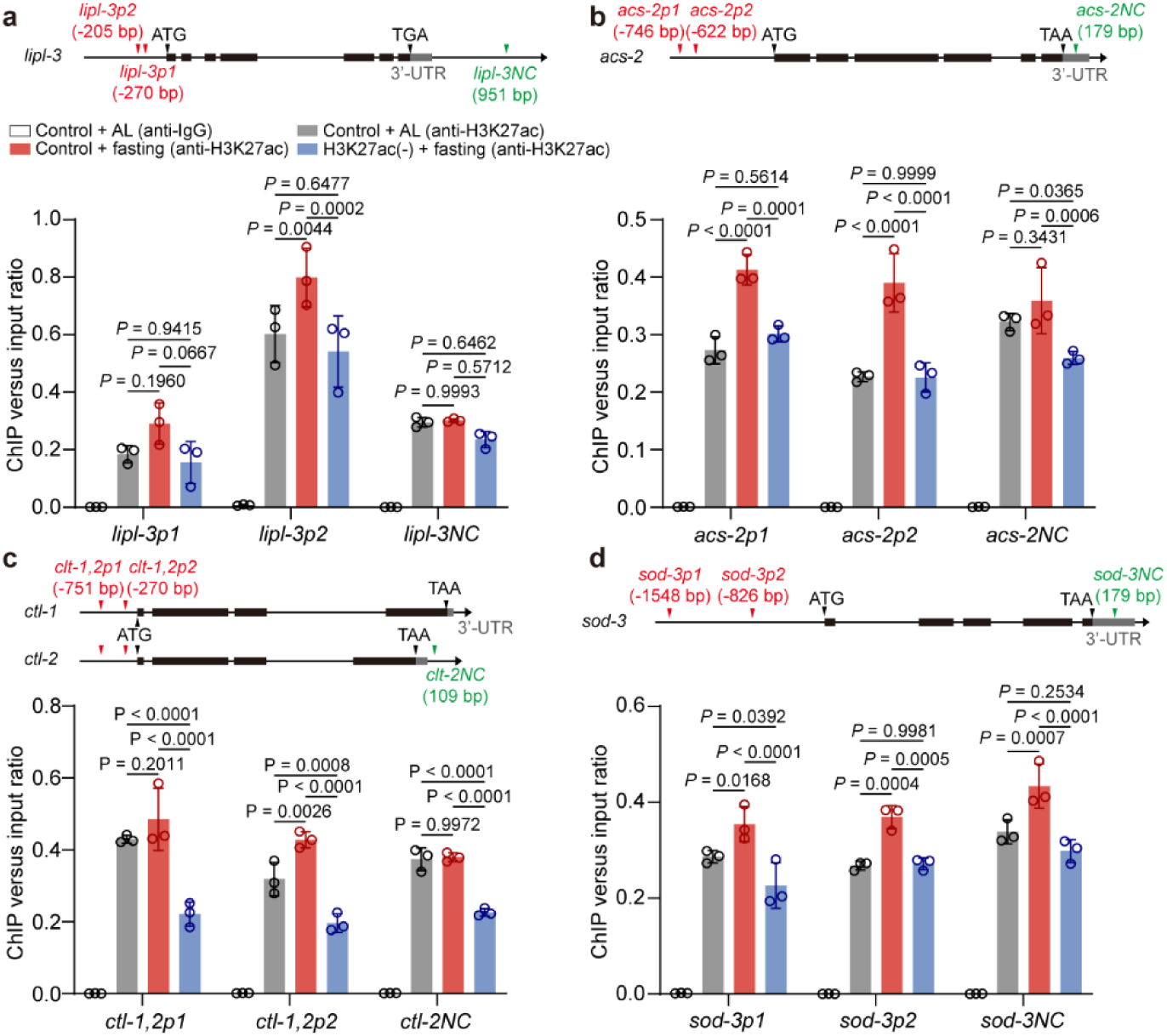
Hormetic fasting elevates H3K27ac on the promoter of lipid catabolic and antioxidant genes. **a**,**b**, ChIP-qPCR analyses reveal that H3K27ac is enhanced at the promoters of *lipl-3* (*lipl-3p*) (**a**) and *acs-2* (*acs-2p*) (**b**), which is blocked in H3K27ac(-) worms. n=3 independent experiments. **c**,**d**, ChIP-qPCR analyses reveal that H3K27ac is elevated at the promoters of *ctl-1,2* (*ctl-1,2p*) (**c**) and *sod-3* (*sod-3p*) (**d**) in a H3K27ac-dependent manner. The sequences of *ctl-1* and *ctl-2* (*ctl-1,2*) promoters are identical, which can be amplified with the same PCR primers. n=3 independent experiments. The flanking sequences at or near the 3’-UTR serve as the negative controls (*lipl-3NC*, *acs-2NC, ctl-2NC, and sod-3NC*) for ChIP-qPCR. Error bars represent SD. Statistical significance was calculated by two-way ANOVA.

### Hormetic fasting activates antioxidant responses to eliminate the accumulated ROS with aging

Since the key enzymes that convert highly active ROS to water are significantly induced by hormetic fasting in a H3K27ac-dependent manner (Fig. 3g,k), we hypothesized that hormetic fasting may enhance resistance to oxidative stress. To validate this hypothesis, we first quantified ROS through the kinetics of 2’,7’-dichlorofluorescein (DCF), a fluorescent probe derived from the non-fluorescent compound 2’,7’-dichlorodihydrofluorescein diacetate (H2DCFDA), induced by ROS when incubated with worm lysate^43, 44^. As a positive control and a widely used drug that induces oxidative stress^45^, paraquat treatment (PQ) increased ROS robustly in worms fed AL (Fig. 5a,b). In contrast, the post-fasting worms showed a ROS level comparable to control (Fig. 5b). Consistent with this finding, hormetic fasting increases the survival rate upon chronic oxidative stress caused by 10 mM PQ across the lifespan of *C. elegans* (Fig. 5c). Hormetic fasting also improves the resistance to acute and high-dosage oxidative stress caused by 100 mM PQ treatment after fasting for 24 hours (Fig. 5d), showing that hormetic fasting improves the resistance to oxidative stress. Accumulated ROS is a well-known driver of age-promoted damage to DNA, protein, and organelles and is responsible for physiological and pathological degeneration in aged animals^10^. We therefore wondered about the role of hormetic nutrient stress and H3K27ac in the accumulated oxidative damage during natural aging. By quantifying DCF fluorescence in worm lysates instead of whole worm staining (as staining is confounded by the autofluorescence in aged animals), we found that hormetic fasting reduces ROS levels in aged worms (D8) to that measured in young animals (D3, Fig. 5e,f). In aggregate, hormetic fasting enhances antioxidant defenses and promotes healthy aging in *C. elegans*.

**Fig. 5.**
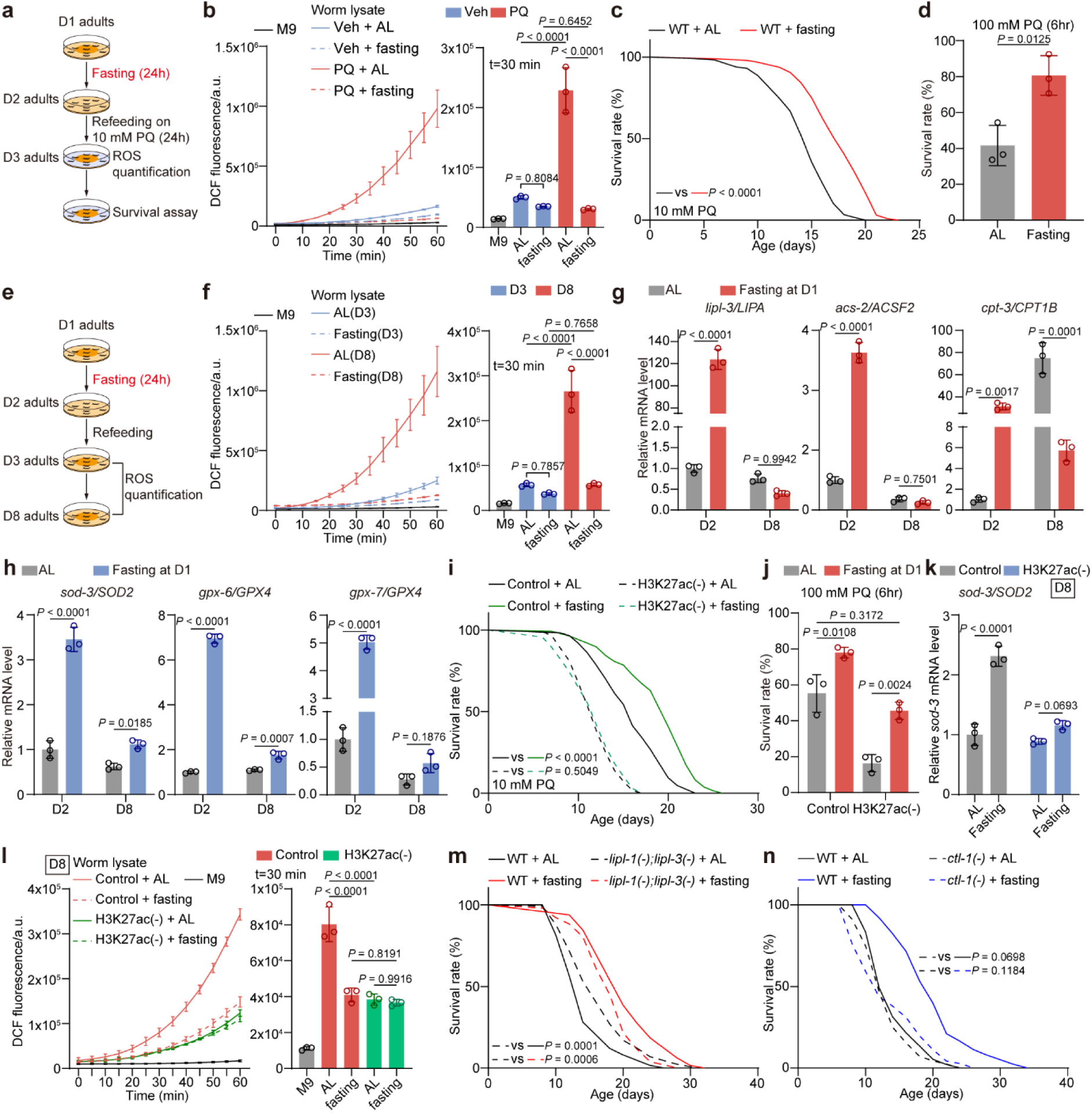
Hormetic nutrient stress mitigates the accumulation of ROS induced by oxidative stress and natural aging. **a**, A depiction of inducing oxidative stress through paraquat (PQ) treatment, followed by ROS assessment and survival analysis assay. **b**, ROS quantified by DCF probe is significantly induced by PQ treatment, which is abolished by fasting at D1. M9 serves as a baseline for the time-course fluorescence quantification. n=3 independent experiments. **c**,**d**, Hormetic fasting for 24 hours at D1 improves the resistance against the chronic (**c**) and acute (**d**) oxidative stress caused by 10 mM and 100 mM PQ treatment from D2, respectively. n=3 independent experiments. **e,** A depiction of assessing ROS in worms at the indicated ages after being subjected to fasting at D1. **f**, ROS quantified by DCF probe is significantly accumulated in aged worms (D8), whereas hormetic fasting at D1 maintains the ROS level compared to that in young worms (D3). M9 serves as a baseline for the time-course fluorescence quantification. n=3 independent experiments. **g**, The lipid catabolic genes are robustly induced by hormetic fasting immediately, whereas their expression is compared with that of AL-fed worms at D8. n=3 independent experiments. **h**, The expression of critical enzymes *sod-3/SOD2* and *gpx-6/GPX4* participating in ROS degradation is persistently elevated in both young and aged worms after fasting at D1 and refed from D2 to D8. n=3 independent experiments. **i**,**j**, H3K27ac inhibition blocks the fasting-promoted resistance against chronic (**i**) and acute (**j**) oxidative stresses, which are caused by 10 mM and 100 mM PQ treatment, respectively. n=3 independent experiments. **k**, Induced *sod-3/SOD2* expression in aged worms by hormetic fasting at D1 is ablated by H3K27ac inhibition. n=3 independent experiments. **l**, H3K27ac inhibition reduced the ROS level in aged worms subject to either AL or fasting. M9 serves as a baseline for the time-course fluorescence quantification. n=3 independent experiments. **m**, *lipl-1(-);lipl-3(-)* mutation partially rescues the lifespan extension caused by hormetic fasting at D1. **n**, *ctl-1(-)* mutation abolishes the lifespan extension promoted by hormetic nutrient stress. See also Supplementary Data 3 for independent biological replicates and summary lifespan statistics. a.u., astronomical unit in (**b**, **f**, and **l**). Error bars represent SD. Statistical significance was calculated by one-way ANOVA (**b**, **f**, **j**, and **l**), unpaired *t*-test (**d**), two-way ANOVA (**g**, **h**, and **k**), and Mantel-Cox Log Rank (**c**, **i**, **m**, and **n**).

Due to the ability of hormetic fasting to reduce ROS levels in old animals to that typical of young worms, we were next interested in the temporal regulation of H3K27ac-dependent gene expression in response to fasting. Intriguingly, by tracking the transcription of lipid catabolic genes in worms subjected to day 1 hormetic fasting across the lifespan, we find that mRNAs are significantly induced after fasting immediately but return to a comparable or lower level than that in worms fed AL by day 8 of adulthood (Fig. 5g), consistent with the activity of energetic stress reporter *acs-2p::GFP* (Extended Data Fig. 5a). In contrast, *sod-3, gpx-6* remain significantly elevated in fasted worms compared to worms fed AL at day 8 (Fig. 5h), indicating a lifelong activation of antioxidant defenses by hormetic fasting. Consistent with the importance of H3K27ac in these expression changes, H3K27ac inhibition also blunts the optimized survival by hormetic fasting under chronic oxidative stress (Fig. 5i) and causes a significant decrease in the resistance against acute and high-dosage oxidative stress (Fig. 5j). Concordantly, H3K27ac inhibition blunts the transactivation of *sod-3* specifically in fasted worms at day 8 (Fig. 5k and Extended Data Fig. 5b), showing that hormetic nutrient stress occurring in young adulthood maintains antioxidant defenses throughout the entire lifespan in a manner completely dependent upon modulators of H3K27 acetylation.

We next asked whether H3K27ac inhibition has an epistatic effect on ROS metabolism in aged worms as well. Unexpectedly, we found that the ROS level is significantly reduced in H3K27ac(-) worms at day 8, regardless of whether they undergo fasting at the young stage (Fig. 5l), which contrasts with the reduced resistance to oxidative stress in H3K27ac(-) worms (Fig. 3k and 5h). Since fatty acid oxidation is hyperactivated by energetic stress to utilize alternative fuels, and serves as the primary source of mitochondrial ROS production^46^, we suspected that the reduced ROS in H3K27ac(-) worms may be attributed to silenced fatty acid oxidation. To test this hypothesis, we utilized a double mutation of lysosomal lipase *lipl-1* and *lipl-3* (*lipl-1(-);lipl-3(-)*) to block lipid catabolism activated by fasting. As expected, *lipl-1(-);lipl-3(-)* double mutation leads to an unchanged fat mass in response to food deprivation (Extended Data Fig. 5c,d). In addition, ROS level is further decreased by *lipl-1(-);lipl-3(-)* double mutation in parallel to the effects from fasting (Extended Data Fig. 5e), indicating that fatty acid degradation is the dominant source of ROS in both aged and fasted worms. *lipl-1(-);lipl-3(-)* mutants also show a mildly longer lifespan compared to WT worms and have a further extended lifespan upon hormetic nutrient stress at day 1 (Fig. 5m). *acs-2* and *cpt-3* knockdown by RNAi partially inhibits the lifespan extension upon hormetic fasting (Extended Data Fig. 5f,g), suggesting a partly dispensable role of lipid catabolism in the longevity prompted by hormetic fasting. On the other hand, the pro-longevity effect of hormetic nutrient stress is blunted by *clt-1(-)* mutation, and significantly suppressed by *sod-3(-)* mutation, *clt-2(-)* mutation, and RNAi knockdown of *gpx-6* and *gpx-7* (Fig. 5n and Extended Data Fig. 5h-k), demonstrating that antioxidant defense is essential for the lifespan extension following exposure to nutrient stress. Adaptive lipid catabolism may thus be more critical for alternative fuel utilization and survival in response to acute environmental challenges.

### Multiple transcription factors are coordinated for the lifespan extension of hormetic nutrient stress

We subsequently directed our attention towards identifying the transcription factors responsible for initiating lipid catabolism and antioxidation in response to nutrient stress. Critical genes in lipid catabolism, such as *lipl-3*, *acs-2*, *cpt-3*, are jointly controlled by NHR-49/HNF4 and TFEB/HLH-30 in longevity paradigms^6, 21, 34^. DAF-16/FOXO is well-known as a transcription factor for antioxidant responses, positively influencing the transactivation of *sod-3*, *ctl-1*, *ctl-2*, *gpx-6*, and *gpx-7* ^47, 48^. We therefore investigated the roles of these transcriptional factors in the adaptive response upon fasting. By qRT-PCR, we found that both elevated lipid catabolic and antioxidant genes upon fasting are blunted by *nhr-49(-)* mutation, and *hlh-30(-)* mutation prevents fasting-induced increases of *lipl-3*, and partially suppresses the induction of antioxidative genes (Fig. 6a). Consistent with its pivotal role in antioxidation, *daf-16(-)* mutation has less impact on lipid catabolism but abolishes the induction of antioxidant genes (Fig. 6a). These results demonstrate that HLH-30 and DAF-16 are primarily responsible for fatty acid and ROS scavenging, respectively, and NHR-49 is the dominant transcription factor that activates both metabolic rewiring and antioxidation (Fig. 6b). This cooperation also suggests that a transcription factor network drives adaptive responses to nutrient stress.

**Fig. 6.**
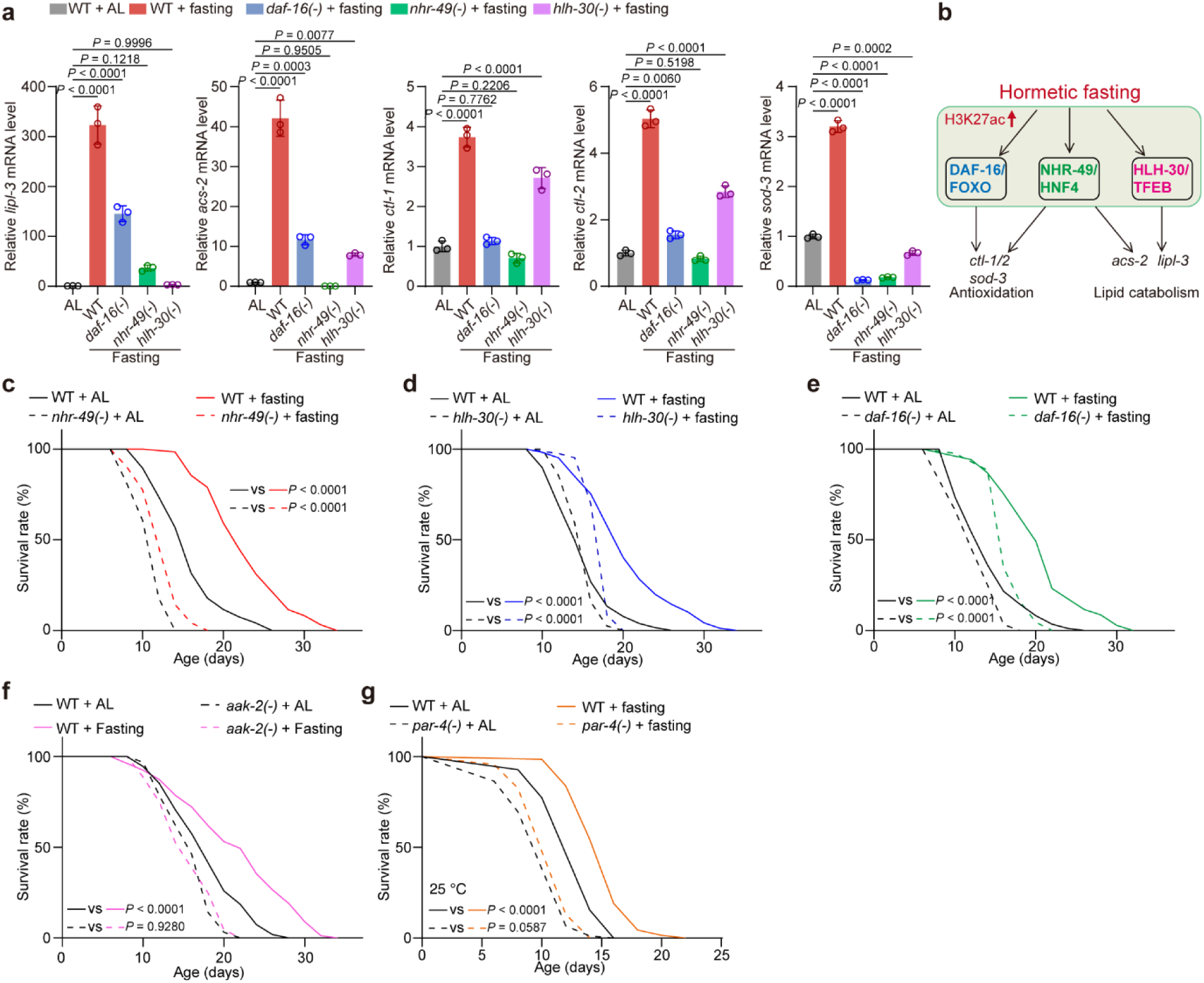
Hormetic fasting orchestrates the transactivation of lipid catabolism and antioxidant defense by employing NHR-49, DAF-16 and HLH-30. **a**, qPCR-PCR of indicated genes in WT worms and indicated mutants when worms are fed ad libitum (AL) or subjected to hormetic fasting at D1. n=3 independent experiments. **b**, A depiction summarizes the result in (**a**) that NHR-49 is necessary for the transactivation of both lipid catabolic and antioxidant genes in response to fasting, whereas DAF-16 and HLH-30 make a more significant contribution to antioxidative and lipid catabolic genes, respectively. **c-e**, Individual mutation of *nhr-49* (**c**), *hlh-30* (**d**), and *daf-16* (**e**) partially rescues the lifespan extension upon hormetic fasting. **f**, *aak-2(-)* mutation abolishes the lifespan extension of hormetic fasting. **g**, A temperature-sensitive mutation of *par-4*/LKB1 blunts the lifespan extension in response to fasting. The survival rate is assessed at 25 °C. See also Supplementary Data 3 for independent biological replicates and summary lifespan statistics. Error bars represent SD. Statistical significance was calculated by one-way ANOVA (**a**) and Mantel-Cox Log Rank (**c-g**).

To our surprise, individual mutations of *nhr-49*, *hlh-30*, or *daf-16* are ineffective in blunting the lifespan extension upon hormetic fasting (Fig. 6c-e), suggesting that these transcription factors contribute to the lifespan extension of stress hormesis collaboratively and/or at least partially redundantly by promoting lipid catabolism and antioxidative response. As the hub of energy sensing and essential regulator for metabolic adaptation, AMPK is activated after food deprivation for 24 hours with elevated phosphorylation level (Extended Data Fig. 6a). A null mutation of *aak-2* (*aak-2(-)*), one of the two orthologues of AMPKα in nematodes, abolishes the lifespan extension upon hormetic fasting (Fig. 6f). A temperature-sensitive loss-of-function mutation of *par-4* (*par-4(-)*), encoding the orthologue of LKB1 that promotes AMPK phosphorylation and activation^49^ (Extended Data Fig. 6b), is also epistatic to the longevity in worms subjected to hormetic nutrient stress (Fig. 6g), demonstrating that LKB1/AMPK axis is the dominant upstream of all the factors promoting longevity upon fasting, which is complied with its actin in activating NUP50 post-translationally^6^.

Mechanistically, the nuclear translocation of HLH-30 and DAF-16 is significantly enhanced by short-term food deprivation and remains unaffected by H3K27ac(-). Nuclear localization of NHR-49 is mildly but insignificantly induced by fasting, and remains unchanged when H3K27ac is inhibited (Extended Data Fig. 7a-c). These data suggest that H3K27ac modulators may regulate the adaptive transcriptome by directly affecting chromatin accessibility for transcription factor binding.

### Enhanced H3K27ac is determined by pioneer factor PHA-4/FOXA and cooperating transcription factor NHR-49/HNF4

As a marker of open chromatin and enhancer-activated transcription, elevated H3K27ac maintains chromatin accessibility and promotes the recruitment of transcriptional modulators. Mechanistically, histone acetyltransferase could bind to target promoters by its intrinsic bromodomains or be recruited by pioneer transcription factors^50^. To determine which of these mechanisms are operative in hormetic fasting, we examined the reciprocal dependency of H3K27ac marks and transcription factors binding on active promoters during fasting. Although the NHR-49 binding on *acs-2* promoter is enhanced by fasting, it is not blocked in fasted H3K27ac(-) worms (Fig. 7a). In contrast, the enhanced H3K27ac on *acs-2* promoter is abolished by *nhr-49(-)* mutation (Fig. 7b), suggesting that NHR-49 binding is necessary for and prior to H3K27ac elevation upon stress. Combined with our previous finding that NHR-49 determines the interaction of NUP50 with downstream promoters^6^, our cumulative evidence suggests that NHR-49 may determine the site-specific H3K27ac upon hormetic fasting. In mammals, NHR-49/HNF4 serves as a cooperating transcription factor for the pioneer factor PHA-4/FOXOA to promote chromatin accessibility prior to recruiting H3K27ac promoting machinery^51^. Since *acs-2* is a shared binding target of PHA-4 and NHR-49 during starvation in worms^21, 52^, we then speculated that PHA-4/FOXA and NHR-49/HNF4 may trigger epigenetic modifications and dictate site-specificity for transcriptional reprogramming during fasting. To deplete PHA-4 efficiently during fasting and avoid negative developmental pleiotropy associated with its loss during development^53, 54^, we constructed an endogenously CRISPR-tagged PHA-4::mKate2::AID::3xFLAG worm strain. Similarly to NHR-49, PHA-4 degradation suppresses the H3K27ac enrichment during fasting (Fig. 7c), showing that the pioneer factor PHA-4 and cooperating factor NHR-49 are both required for recruiting H3K27ac machinery for the subsequent transactivation of lipid catabolism and antioxidation. Furthermore, H3K27ac(-) barely affects the interaction of NPP-16 and the *acs-2* promoter, but NPP-16 is essential for the H3K27ac enrichment on *acs-2* (Fig. 7d,e). In aggregate, despite stability of global H3K27ac levels during nutrient stress, H3K27ac is specifically elevated at the promoter of the lipid catabolic genes in a manner dependent on the pioneer factor PHA-4, the cooperating factor NHR-49, and the chromatin-associated nucleoprotein NPP-16/NUP50 (Fig. 7f).

**Fig. 7.**
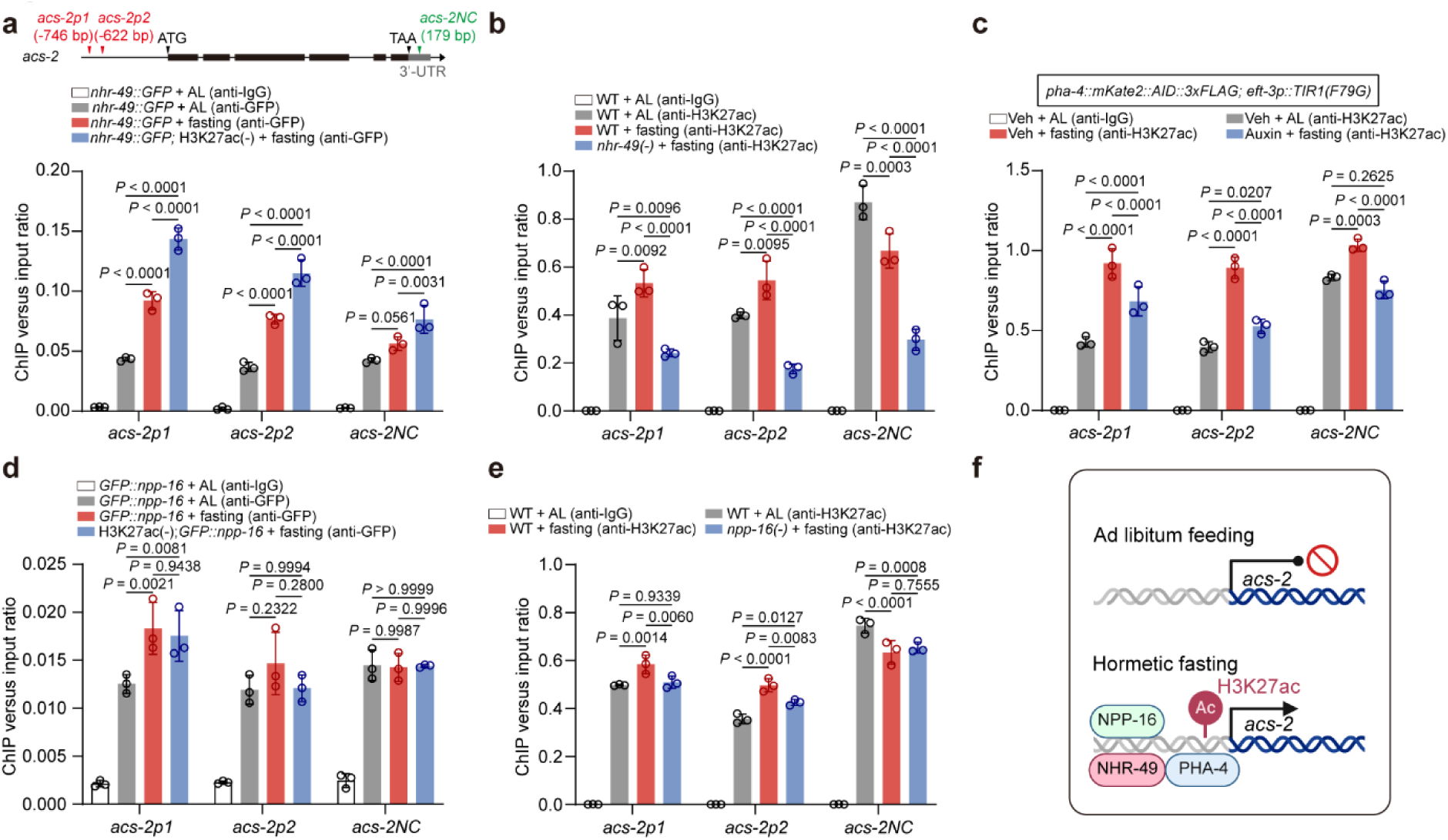
**The H3K27ac enhancement on active promoter is determined by pioneer transcription factors and NPP-16. a-c**, ChIP-qPCR analyses reveal that H3K27ac modulators are dispensable for the interaction between NHR-49 and the *acs-2* promoter (**a**), whereas *nhr-49(-)* mutation (**b**) and PHA-4 degradation (**c**) prevents the H3K27ac induction on *acs-2* promoter in response to hormetic fasting. n=3 independent experiments. **d**,**e**, The fasting-promoted chromatin association of NPP-16 is unchanged in H3K27ac(-) worms (**d**), but NPP-16 is necessary for the H3K27ac enrichment on the *acs-2* promoter in fasted worms (**e**). n=3 independent experiments. **f**, A depiction summarizing the essential role of NHR-49/HNF4, PHA-4/FOXA, and NPP-16/NUP50 in elevating H3K27ac at the promoter of lipid catabolic gene. Error bars represent SD. Statistical significance was calculated by two-way ANOVA.

## Discussion

CR is among the most influential and practical longevity paradigms, known to reduce aging-related disorders across a wide range of species^2–4^. However, in addition to lifelong loss of lean mass and changes in immune repertoire^2, 55^, a common adverse effect observed across most CR regimens is delayed development primarily due to reduced energy availability^56^.

Similarly, other longevity paradigms associated with energy sensing, such as insulin/IGF-1 Signaling (IIS) inhibition, ETC inhibition, and biguanide treatment, also have delayed developmental timing^56, 57^. Here, we describe an adulthood-onset nutrient stress that leads to lifespan extension without altering development. By RNAi screen and transcriptomic analysis, this study establishes H3K27ac as a transcriptional switch that maintains redox homeostasis by coordinating lipid catabolism and antioxidant defense in response to hormetic nutrient deprivation, thereby promoting healthy aging and longevity in *C. elegans*.

Building on our previous research^6^, we demonstrate an AMPK-NUP50-H3K27ac axis that relays energy sensing to transcriptional and metabolic reprogramming for longevity, establishing a heretofore unappreciated molecular basis for hormetic nutrient stress in healthy aging (Fig. 8).

**Fig. 8.**
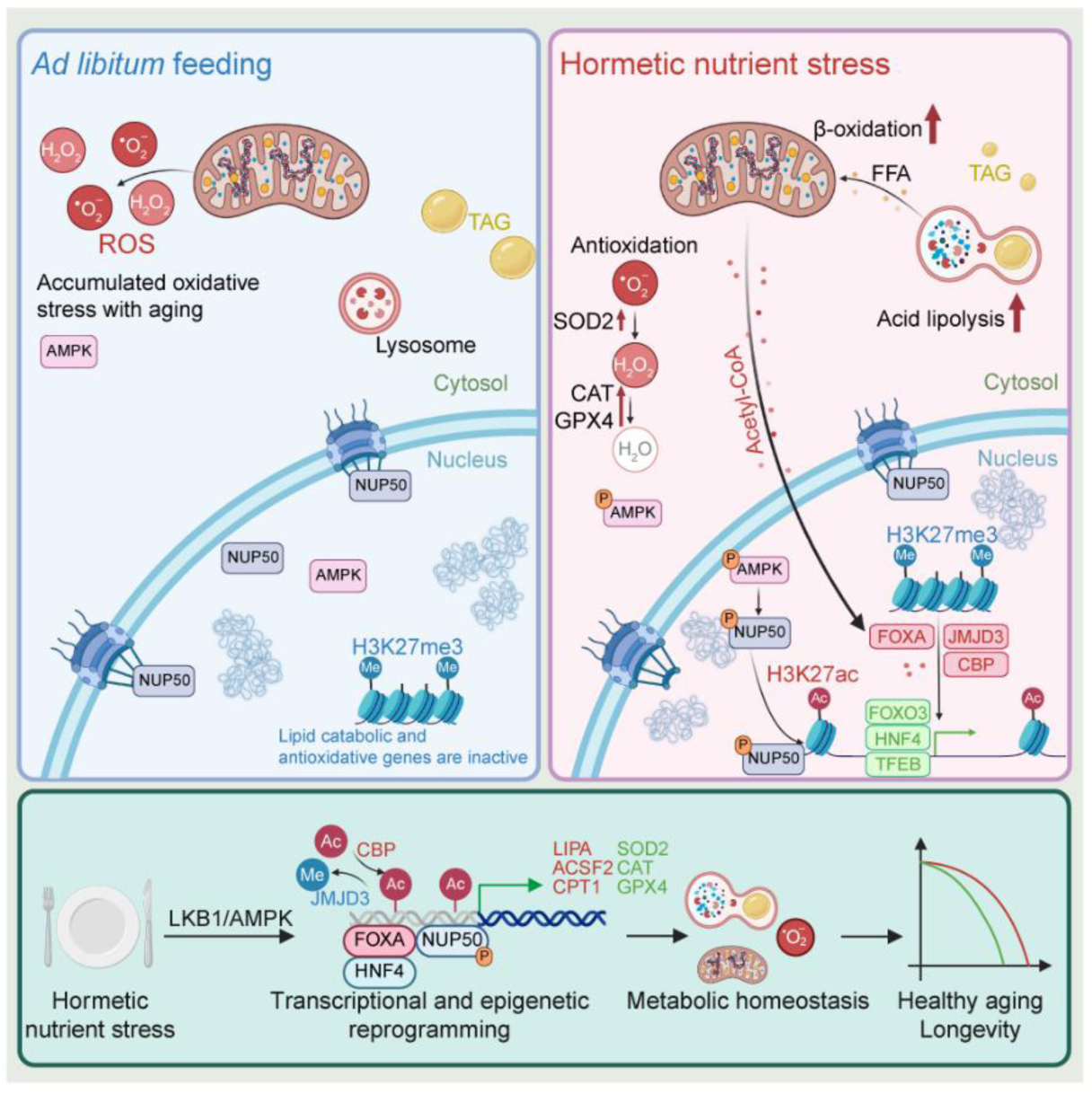
**A schematic illustration of H3K27ac facilitating transcriptional and epigenetic reprogramming to maintain metabolic homeostasis and promote longevity in response to hormetic nutrient stress.**

CR and CR-like regimens in worms are induced in diverse ways to promote longevity, such as reducing food intake through diluted bacterial feeding (sDR) and genetic mutation of *eat-2* that slows down the pharyngeal pumping rate^56^. Biguanide treatment also leads to CR-like phenotypes in metabolic rewiring and longevity^58^. Imminent fasting from adulthood is sufficient to promote longevity in *C.elegans*^59^. In combination with our previous study^6^, we described that adulthood-onset fasting for 24 hours, without any perturbation to development, is effective in extending lifespan by coordinating lipid catabolism and antioxidant defenses, providing new insights into longevity paradigms associated with nutrient stress. Although most CR and CR-like paradigms share a positive effect on healthy aging and promoting longevity, distinct paradigms have discreet effector mechanisms. As a metabolic hub for energy sensing and adaptive regulator, AMPK is necessary for the longevity caused by sDR but is dispensable for the longevity caused by *eat-2(-)* mutation^5^. In this study, we demonstrated that the LKB1/AMPK axis is essential for the longevity of hormetic nutrient stress, in accordance with our previous finding that AMPK-induced NUP50 is critical for the metabolic rewiring and longevity upon nutrient stress. In addition, we also identified H3K27ac as the transcriptional switch that activates lipid catabolism immediately after fasting and antioxidant defenses persistently across the lifespan. These findings illustrate a clear pathway connecting energy sensing to epigenetic rewriting and transcriptional reprogramming, thereby promoting longevity through maintaining redox homeostasis.

As reported^6^, NUP50 is induced post-translationally by AMPK in response to nutrient stress, serving a critical role in the longevity paradigms associated with energetic stress by mediating chromatin plasticity. The transcription factor NHR-49/HNF4 governs the association of NUP50 with chromatin to facilitate transcriptional activation. In this study, we identified H3K27ac as the factor linking NUP50 activation and chromatin remodeling, thereby enhancing chromatin accessibility in response to nutrient stress. Although alterations in H3K27ac have been reported in response to various stressors, the majority of these changes occur at the level of global H3K27 methylation or acetylation. This study demonstrates that the site-specific enrichment of H3K27ac is regulated by the pioneer transcription factor and cooperating factors, revealing a conserved mechanism within the FOXA-HNF4-H3K27ac axis in chromatin remodeling in murine liver^51, 60^. Additionally, we identify nucleoporin NUP50 as a crucial regulator within this pathway, responsible for positioning H3K27ac on the worm genome. NUP50 also plays a key role in promoting longevity in the nematode intestine^6^, which is the most critical metabolic tissue, functionally analogous to the liver. These findings deepen the understanding of the FOXA-HNF4-H3K27ac axis in physiological responses to stress in adulthood, extending beyond its role in development and cell fate determination.

Nutrient availability governs metabolic homeostasis and determines the preferential substrate used in catabolism. In response to food deprivation, fatty acid oxidation becomes the primary energy source in place of glycolysis. However, fatty acid oxidation accelerates ROS generation by enhancing the TCA cycle. Adaptively, SOD2 is activated for antioxidation when animals are subjected to dietary restriction^12^. Here, we uncovered a new adaptive program that eliminates redundant ROS upon hormetic fasting, illustrating an H3K27ac-dependent transcriptome that broadly transactivates antioxidant genes involved in ROS scavenging, including superoxide dismutase, catalase, and glutathione peroxidase. Intriguingly, this adaptive antioxidant defense persistently maintains ROS at a low level across different stages of life, leading to improved stress resistance and lifespan extension.

In summary, we identify pioneer factors guided H3K27ac as the key transcriptional switch that activates lipid catabolism for alternative energy fueling and promotes antioxidant defense in response to nutrient stress, thereby eliminating the ROS generated from fatty acid oxidation and accumulated in aged animals. Furthermore, complementing our previous research^6^, we demonstrated an unappreciated mechanism that transmits the LKB1/AMPK pathway to the NUP50/H3K27ac axis, thereby facilitating epigenetic and transcriptional reprogramming. This work links energy sensing to defensive responses for hormesis and underpins the beneficial effects of hormetic fasting on longevity promotion.

## Methods

### *C. elegans* genetics and maintenance

*C. elegans* were grown on the normal growth medium (NGM) supplemented with 0.1875 mg/ml streptomycin and fed with the E.coli strain OP50-1 or HT115(DE3) with indicated RNAi under standard conditions at 20 °C unless otherwise noted^61^. *par-4(it57)* mutants were grown at 15°C, and moved into 25 °C to induced par-4 depletion from hatching. All the strains used in this paper are listed in Supplementary Data 5. Some of them were provided by the Caenorhabditis Genetics Center funded by the NIH Office of Research Infrastructure Programs (P40 OD010440), National BioResource Project (NBRP), the laboratory of William Mair at the Harvard T.H. Chan School of Public Health, and the laboratory of Javier Irazoqui at UMass Chan Medical School.

For auxin feeding, the auxin analog 5-Ph-IAA (Bioacademia, Cat#30-003) was dissolved in DMSO to make a stock of 100 mM and diluted to 100 μM in water as a working solution freshly before using. Thereafter, 450 μl of the working solution was added to the 9 mL of NGM in a 6 cm petri dish seeded with OP50 or HT115 bacteria, leading to a final concentration of 5 μM in agar. Auxin was added to plates one day before plate use, and plates were kept in the dark. The DMSO of the same dilution rate serves as the negative control.

For hormetic fasting, worms synchronized by bleaching and L1 arrest were grown on seeded NGM or RNAi plates with abundant food until D1 adulthood, then transferred to unseeded NGM or RNAi plates for 24 hours of fasting treatment. The fasted worms were subjected to the following assessment immediately after 24-hour fasting unless otherwise noted.

### Plasmid construction

To generate pJW1586-*pha-4::mKate2::AID::3xFLAG* as the repair template for *pha-4* genomic editing, 572 bp of upstream and 542 bp downstream sequence flanking the stop codon of genomic *pha-4* were cloned into the pJW1586 vector digested by SpeI and AvrII through Gibson assembly. The pJW1586 vectors were provided by Addgene (Cat#121057).

To generate plasmids expressing guide RNAs for CRISPR editing, pDD162-*npp-16* and pDD162-*npp-pha-4* the sgRNA sequences were designed by CRISPOR (http://crispor.gi.ucsc.edu/crispor.py)^62^. The guide sequence (GGCGGCCGAGTTCGGGTTGG) targeting the C-terminus of *pha-4* was cloned into pDD162 by PCR and T4 DNA ligase (New England Biolabs, Cat#M0202S). The pDD162 vector was provided by Addgene (Cat#47549).

### *C. elegans* strain generation

For the genome editing of *pha-4* by CRISPR/Cas9, plasmids of pDD162-*npp-16*(10 ng/μl) and pJW1586-*pha-4::mKate2::AID::3xFLAG* (50 ng/μl) were co-injected into N2 worms with an injection marker of *myo-2::mCherry* (2.5 ng/μl). Salmon sperm DNA was added as a carrier to bring the injection mix final concentration to 100 ng/μl of DNA. The homozygotes with a single copy knock-in were screened with the hygromycin resistance assay as reported^63^.

### RNA interference

RNAi clones were isolated and sequence validated from the Ahringer library^20^. RNAi plates were made with the standard NGM recipe supplemented with 5 mM IPTG and 200 μg/ml carbenicillin. After overnight culture in LB containing 200 μg/ml carbenicillin at 37 °C, the bacteria were concentrated 5-fold by centrifugation. All RNAi experiments were conducted from hatching unless otherwise noted.

For the epigenetic library RNAi screen, about 15,000 synchronized WT worms were grown to L4 on the RNAi medium seeded with HT115 carrying empty vector in three 10 cm dishes and transferred onto 24-well plates seeded with HT115 carrying the epigenetic RNAi library containing 50 mM 5-fluoro-2′-deoxyuridine (FUdR) after washing with M9 for three times to remove the bacteria as much as possible. The worm concentration in each well is approximately 40 worms/well. For RNAi library preparation, the indicated RNAi clones were isolated from the Ahringer library^20^, and cultured in 1.5 mL LB containing 200 μg/ml carbenicillin on the deep well plates at 37 °C overnight. The bacterial liquid was centrifuged at 4000 g for 10 minutes and suspended in 40 μl LB with carbenicillin. The indicated bacteria were seeded on each well containing 0.5 mL RNAi medium in 24-well plates. The survival rate is quantified at day 20 of adulthood. Two independent experiments were performed, and the final ranking was calculated according to the average survival rate of the two biological replicates. The raw statistics of the RNAi screen are shown in Supplementary Data 1.

### Microscopy

Microscopic images were taken on a Leica DM6 B microscope with THUNDER Imager. For imaging of transcription reporters, about 15 worms at the indicated ages were mounted on 5% agar pads and anesthetized by 10 mM sodium azide and imaged on a Leica DM6 B microscope with Thunder Imager and a 5x objective. For imaging of the nuclear localization of transcription factors, 10-15 worms were mounted on 5% agar pads and anesthetized using 5 mM levamisole (Sigma-Aldrich, Cat#L9756), and imaged on a Leica DM6 B microscope with Thunder Imager and a x63 objective for each biological replicate. To image Nile Red stained worms, the worm pellet was dropped onto the slice and subjected to imaging on a Leica DM6 B microscope with Thunder Imager with a 10x objective.

### Image analysis

For the quantification of transcription reporters, about 15 worms were laid on the agar pad side by side, and the mean intensity of the entire worm area was quantified by Image J. For Nile Red quantification, the posterior intestinal cells from 10-15 worms were selected and quantified using Image J for each biological replicate. For the nuclear localization of transcription factors, the GFP intensity in nuclei and cytoplasm of the posterior intestinal cells was quantified by Image J. All fluorescent quantifications were normalized to mean of the control group.

### Lifespan assay

Worms were synchronized through the overnight egg laying. After approximately 60 hours, about 40 late L4s were picked onto each 6 cm NGM agar plates seeded with OP50 or HT115, containing 50 mM 5-fluoro-2′-deoxyuridine (FUdR). Every biological replicate is performed on five 6 cm NGM agar plates containing about 200 worms. Worms were counted every other day, and the ones not exhibiting spontaneous movement or subsequently not responding to mechanical prodding were scored as dead. Animals that exhibited bursting vulva or plates that became contaminated were censored. Statistical analysis was performed using the Mantel-Cox Log Rank in GraphPad Prism. See also Supplementary Data 3 for independent biological replicates and summary lifespan statistics.

### Nile Red staining

Neutral lipids were stained and quantified by Nile Red staining as described^64^. Briefly, Nile Red (Fisher Scientific, Cat# N1142) was dissolved in acetone to a stock solution concentration of 5 mg/ml and stored in the dark. Before staining, the working solution was freshly diluted to 30 μg/ml by 40% isopropanol. Approximately 200 young worms at indicated ages were synchronized through egg-laying and subsequently collected for staining. After 3x washing by PBST (1x PBS with 0.01% Triton X-100) to remove as much bacteria as possible, the worms were fixed by incubating with 100 μl 40% isopropanol for 3 min at room temperature. The fixed worms were stained in 400 μl Nile Red working solution by rotating for 2 hours at room temperature, then centrifuged and suspended in 400 μl 1x PBST. The worm pellet was subject to imaging by a Leica DM6 B microscope with THUNDER Imager after rotating in PBST for 30 min.

### Chromatin immunoprecipitation (ChIP)

ChIP was performed as reported with minor modifications^65^. In brief, about 50000 worms synchronized by bleaching and L1 arrest were grown to L4 on NGM or RNAi medium seeded with OP50 or the indicated RNAi bacteria, respectively. For the ChIP in npp-16OE worms, worms were collected on D1 when they had just reached adulthood. For the ChIP experiments in fasted worms. L4 worms were transferred to the indicated medium containing 50 mM FUdR and fed bacteria, and moved to unseeded plates at D1 for fasting, and collected after 24 hours of food deprivation. After washing with M9 three times to remove bacteria as much as possible, worms were crushed in homogenizers and crosslinked with 1% formaldehyde (vol/vol) in PBS for 20 min at room temperature. The crosslinking was then quenched by adding 200 μl 2.5 M glycine and incubation for another 20 min. After washing three times with PBS containing protease inhibitor cocktail (Sigma-Aldrich, Cat# 11873580001), the worm pellet was suspended in 1 ml lysis buffer (50 mM HEPES-KOH, pH 7.5, 150 mM NaCl, 1 mM EDTA, 0.1% (wt/vol) sodium deoxycholate, 1% (vol/vol) Triton X-100, 0.1% (wt/vol) SDS, protease inhibitor cocktail). The lysate was split into 500 μl aliquots in 1.5 mL tubes and then sonicated using a Qsonica Q800R Sonicator with a power setting of 70% and a 30-second on/30-second off pulse for a total of 20 minutes of sonication time. The protein supernatant was isolated by centrifugation at 1,6000 g 4°C for 15 min, and then adjusted to 20 mg/ml with lysis buffer after being quantified with the BCA kit (Thermo Fisher Scientific, Cat# 23225). The supernatant was split into 1 ml for each reaction, and 50 μl of it was kept at -80 °C as the input sample. The antibody was added to 1 ml (2 mg protein) supernatant and incubated at 4 °C overnight. 5 μg anti-GFP (Cat# A-11120, Thermo Fisher Scientific) or anti-H3K27ac (Cat# ab4729, Abcam) antibody was added for each reaction. After the antibody incubation, 50 μl slurry of Protein A/G magnetic agarose beads (Thermo Fisher Scientific, Cat# 78610) was added after pre-washing with lysis buffer three times in order to capture the antibody and target protein, and incubated for 1 hour at 4 °C. The magnetic beads were washed twice with W1 buffer (50 mM HEPES-KOH, pH 7.5, 150 mM NaCl, 1 mM EDTA pH 8.0, 1% (wt/vol) sodium deoxycholate, 1% (vol/vol) Triton X-100, 0.1% (wt/vol) SDS, and protease inhibitor cocktail), twice with W2 buffer (50 mM HEPES-KOH, pH 7.5, 1 M NaCl, 1 mM EDTA, pH 8.0, 0.1% (wt/vol) sodium deoxycholate, 1% (vol/vol) Triton X-100, 0.1% (wt/vol) SDS, and protease inhibitor cocktail), once with W3 buffer (50 mM Tris-Cl, pH 8.0, 0.25 mM LiCl, 1 mM EDTA, 0.5% (vol/vol) NP-40 and 0.5% (wt/vol) sodium deoxycholate), and three times with 1xTE (10 mM Tris-Cl, pH 8.0, 1 mM EDTA in water). The protein on the magnetic beads was then digested with 10 μg proteinase K (Thermo Fisher Scientific, Cat# 26160) at 45 °C for 2 hours, and the protein in 50 μl input was digested by 10 μg proteinase K at 55 °C for 4 hours. Cross-links were reversed by incubation at 65 °C overnight, releasing the DNA fragments. The DNA in each sample was extracted twice with 250 μl phenol–chloroform–isoamyl alcohol (Sigma-Aldrich, Cat# P3803) and centrifuged at 14,000 g for 10 minutes at room temperature. The DNA was then pelleted in 1 ml ethanol containing 1 μg glycogen (Thermo Fisher Scientific, Cat# R0551) with incubation at -80 °C for 20 minutes and centrifuged at 14,000 g for 10 minutes. After washing with 500 μl 70% ethanol, the DNA pellet was air-dried briefly, and dissolved in 200 and 40 μl elution buffer (10 μM Tris-HCl, pH 8.5) for input and ChIP samples, respectively. For downstream reactions, the DNA was 10x diluted with water and used as the template in subsequent PCR reactions.

### qRT-PCR

Approximately 1000 worms were synchronized by bleaching and L1 arrest and then collected into 500 μl RNAzol (Sigma-Aldrich, Cat# R4533) after washing three times with M9. The worms were crushed with a TissueLyser II (QIAGEN) at a frequency of 25 times/sec for 2 minutes in 500 μl RNAzol and 200 μl RNase-free water. After incubating at room temperature for 15 min, the samples were centrifuged at 12,000 g for 15 min. The supernatant was added to an equal volume of isopropanol (about 680 ml) and mixed by vortexing. After incubating at room temperature for 15 minutes, the RNA was pelleted by centrifuging at 12,000 g for 10 minutes. The RNA pellet was air-dried after twice washing with 400 ml 70% ethanol and finally dissolved in 30 μl RNase-free water. The RNA concentration and quality were then determined by NanoDrop One C Microvolume UV-Vis Spectrophotometer (Thermo Fisher Scientific). The cDNA was generated by QuantiTect Reverse Transcription Kit (QIAGEN, Cat# 205314) following the manufacturer’s instructions. The cDNA product from 500 ng of input RNA was diluted 500-fold with DNase/RNase-free water as the template for qPCR. The qPCR was performed with QuantiTect® SYBR® Green PCR (QIAGEN, Cat# 204145) following the manufacturer’s instructions on the CFX96 Real-Time System (Bio-Rad). *act-1* mRNA serves as the internal control for quantifying the corresponding cDNA. Primers used for qPCR are listed in Supplementary Data 6.

### RNA sequencing

All RNA-sequencing data pro-processing, quantification, and statistical testing was performed as previously described^6^. Briefly, ∼1000 synchronized D1 WT and H3K27ac(-) nematodes were fed *ad libitum* or fasted for 24 hours and subsequently collected into 500 μl RNAzol (Sigma-Aldrich, Cat# R4533) after 3x washing for total RNA extraction in biologically independent quadruplicates (16 total samples). The total RNA was extracted, and the genomic DNA was removed using the Direct-zol RNA Miniprep Plus Kit (ZYMO research, Cat# R2072) following the manufacturer’s instructions. The extracted total RNA was evaluated for quality control using a NanoDrop One C Microvolume UV-Vis Spectrophotometer (Thermo Fisher Scientific), with an A260/A280 > 1.98 and an A260/A230 > 1.85. Samples were sent to Azenta (Genewiz) for additional quality control, library preparation, and mRNA sequencing. Samples were first validated for RNA integrity with a RIN score > 8.5 and DV200 > 50 using an RNA Tapestation 4200 (Agilent). Illumina library preparation was performed using polyA selection for mRNA species. Approximately 17.5 - 30 million paired-end 150bp reads were generated per sample, with > 90% of bases passing a Phred quality score Q30.

Fastq read quality control, adapter trimming, quality score filtering, and quasi-alignment were all performed using custom UNIX/Bash shell scripts on the Mass General Brigham ERISTwo Scientific Computing CentOS 7 Linux Cluster. Reads were analyzed for quality control using FastQC v0.11.8^66^ (http://www.bioinformatics.babraham.ac.uk/projects/fastqc^)^ and MultiQC v1.19^67^. Reads were then filtered for Illumina adapter contamination, truncated short reads or low-quality base calls using BBDuk (BBTools)^68^. The subsequently trimmed and cleaned reads were then quasi-aligned to the Caenorhabditis elegans reference transcriptome annotation (WBcel235, Ensembl Release 105) using Salmon v1.9.0^69, 70^, correcting for GC content and sequencing bias using the command parameters ‘--gcBias’ and ‘--seqBias’. All statistical analyses and visualizations were generated using the R v4.3.2 (www.r-project.org) Bioconductor v3.18 statistical environment on a local machine through Jupyter Notebook v6.4.10 (https://jupyter.org)^71^. Quasi-aligned transcript quantification files for each sample were collapsed into gene-level count matrices using R package tximport v1.30.0^72^. The raw statistics of transcripts per million are shown in Supplementary Data 4.

Paired differential expression was calculated using R package DESeq2 v1.42.1x with a design formula of ‘∼ Genotype_Treatment’^73^. Genes were considered differentially expressed with a Benjamini-Hochberg False Discovery Rate (FDR) corrected *P* value < 0.05 and an absolute log_2_ transformed fold change of 1^74^. Metabolism gene set variation analysis (GSVA) was performed using R package GSVA v1.50.1 using metabolic gene sets extracted from WormPaths^35, 75^. Heatmaps were generated using R package pheatmap v1.0.12, using row-scaled z scores generated from DESeq2 variance stabilized count transformation^73^.

Gene set overrepresentation analyses for genes identified as differentially expressed (with thresholds set as indicated above) were performed using R package clusterProfiler v4.12.3 for GO Molecular Function and Reactome-based pathway annotations^38^. Venn diagram overlaps between differentially expressed genes in this study, NPP-16 overexpression studies^6^, and *cbp-1*/*cco-1* knockdown/target studies^15^ were performed using ggvenn v0.1.10. Volcano plots were generated using ggplot2 v3.5.0. Visualizations were post-edited for font, sizing, and appearance using Adobe Illustrator and the Adobe Creative Cloud Suite. All custom bash scripts, Jupyter Notebook files, and processed fastq and transcript count files used in these analyses will be provided upon reasonable request by the corresponding author.

### ROS quantification

ROS was quantified as the fluorescence of DCF as reported with minor modifications^44, 76^. Approximately 500 synchronized worms at the indicated ages were collected and washed three times in M9 to remove as many bacteria as possible. Remove M9 after washing (worm pellet is about 50 μl) and add 50 μl non-denaturig lysis buffer (20 mM Tris HCl pH 8, 137 mM NaCl, 10% glycerol, 1% Nonidet P-40 (NP-40), 2 mM EDTA, protease inhibitor cocktail).

Worms were crushed by Fisherbrand™ Pellet Pestle in 1.5-ml tubes. The protein supernatant was then isolated by centrifuging at 16,000 rpm, 4 °C for 15 min, and its protein concentration was quantified using a BCA kit (Thermo Fisher Scientific, Cat# 23225). Worm lysate containing 5 μg protein was diluted to 10 μl with lysis buffer in a 96-well plate, and 5 μl M9 plus 5 μl lysis buffer served as a negative control. DCFDA (Thermo Fisher Scientific, Cat# D399) was dissolved by DMSO to a stock solution of 50 mM. 200 μlDCFDA working solution was freshly diluted to 50 μM with water and added to each well of the 96-well plate. The fluorescence of DCF was measured every 5 min for 1 hour by DTX 880 Multimode Detector (Beckman Coulter) at an excitation of 485 nm and emission of 535 nm. Each sample has two technical replicates on the same 96-well plate, and the mean of the fluorescent signal was defined as the result of one biological replicate.

### Immunoblotting

Approximately 5,000 worms were synchronized by bleaching and L1 arrest, then plated and collected at the L4. After washing three times to remove bacteria, the worm pellet was added to 100 μl RIPA buffer (50 mM Tris-HCl pH 7.4, 150 mM NaCl, 1% Triton-100X, 1% sodium deoxycholate, 0.1% SDS, 1 mM EDTA, supplemented with protease inhibitor cocktail). Additional phosphatase inhibitors (0.1mM Na_3_VO_4_, 5mM beta-glycerophosphate, 1μM okadaic acid sodium salt, 10mM NaF, 1mM sodium molybdate) were used for blots against p-AMPK. The worms were crushed with a tissue grinder about 100 times. The protein supernatant was clarified by centrifugation at 14,000 g at 4°C for 15 minutes to remove insoluble debris, and protein concentration was determined by BCA kit (Thermo Fisher Scientific, Cat# 23225). After adding 4x loading buffer (Bio-Rad, Cat# 1610738) containing 10% β-mercaptoethanol, and boiling at 95 °C for 10 minutes, protein in the supernatant was separated by SDS-PAGE (Bio-Rad, Cat# 456-1084) and transferred electrophoretically to nitrocellulose membranes (Bio-Rad, Cat#1620115). Blots were blocked with 5% non-fat milk or BSA (for detection of phosphorylation) in Tris-buffered saline with 0.1% Tween 20 detergent (TBST) to reduce background. The membrane was blotted with primary antibody against H3K27ac (1:5000 dilution, Abcam, Cat# ab4729), Histone H3 (1:5000 dilution, Abcam, Cat# ab1791), p-AMPK^Thr172^ (1:2000 dilution, Cell Signaling Technology, Cat# 50081), FLAG (1:5000 dilution, Sigma-Aldrich, Cat# F1804), and β-actin(1:5000 dilution, Abcam, Cat# ab14128) at 4°C overnight, and subsequently the appropriate secondary antibody, either Goat anti-Rabbit IgG (1:5000 dilution, Thermo Fisher Scientific, Cat# 31462) or Goat anti-Mouse IgG (1:5000 dilution, Thermo Fisher Scientific, Cat# G-21040). The blotting signals were captured with the ChemiDoc™ MP Imaging System (Bio-Rad) and quantified by ImageJ.

### Statistics and reproducibility

The RNA-seq analysis was performed as described above. Other statistical analyses were performed by GraphPad Prism with the indicated methods described in the figure legends. No statistical methods were used to predetermine sample sizes, but our sample sizes are similar to those reported previously^6, 77, 78^. The statistical significance of the lifespan was determined using a two-sided log-rank test, as detailed in the Supplementary Information. Data collection and analysis were performed in a blind manner to the experimental conditions.

## Data availability

This paper does not create any original code. The RNA-seq data have been deposited in the GEO database and assigned to the accession number ‘GSE306611’. The source data are available with this paper. Any additional information required to reanalyze data reported in this paper is available from the lead contact upon request.

## Acknowledgements

We thank Dr. William Mair and Dr. Javier Irazoqui for the kind gift of worm strains. We thank members of Soukas lab for the helpful discussions and critical reading of the manuscript. This work was supported by NIH/NIA grants R01AG058256 and R01AG069677 (to AAS), the Weissman Family MGH Research Scholar Award (to AAS), and a gift from Stuart and Suzanne Steele and the Obesity Research Fund at the Mass General Research Institute Center for Genomic Medicine. This research was conducted in part while Y.Z. was a Glenn Foundation for Medical Research Postdoctoral Fellow. Thank the NIH/NIDDK-funded NORC of Harvard (P30DK040561) and the NIH/NIDDK-funded Boston-Area DERC (P30DK135043) for core services. Some worm strains in this study were provided by the Caenorhabditis Genetics Center (CGC) funded by the NIH Office of Research Infrastructure Programs (P40OD010440).

## Author contributions

Conceptualization, Y.Z. and A.A.S.; Methodology, Y.Z. and A.A.S.; Formal analysis, Y.Z. and F.A.; Investigation, Y.Z., F.A., and S.L.; Resources, Y.Z. and A.A.S.; Writing-Original Draft, Y.Z. and A.A.S; Writing-Review & Editing, Y.Z., F.A., S.L., and A.A.S.; Supervision, A.A.S.; Funding Acquisition, A.A.S.

## Reference

1. Gems, D. & Partridge, L. Stress-response hormesis and aging: “that which does not kill us makes us stronger”. Cell Metab 7, 200–203 (2008).

2. Di Francesco, A. et al. Dietary restriction impacts health and lifespan of genetically diverse mice. Nature 634, 684–692 (2024).

3. Brandhorst, S. et al. Fasting-mimicking diet causes hepatic and blood markers changes indicating reduced biological age and disease risk. Nat Commun 15, 1309 (2024).

4. Longo, V.D., Di Tano, M., Mattson, M.P. & Guidi, N. Intermittent and periodic fasting, longevity and disease. Nat Aging 1, 47–59 (2021).

5. Greer, E.L. & Brunet, A. Different dietary restriction regimens extend lifespan by both independent and overlapping genetic pathways in C. elegans. Aging Cell 8, 113–127 (2009).

6. Zhou, Y., Ahsan, F.M. & Soukas, A.A. The nuclear pore complex connects energy sensing to transcriptional plasticity in longevity. bioRxiv (2025).

7. Bennett, C.F. et al. Activation of the mitochondrial unfolded protein response does not predict longevity in Caenorhabditis elegans. Nat Commun 5, 3483 (2014).

8. Mattson, M.P. The cyclic metabolic switching theory of intermittent fasting. Nat Metab (2025).

9. Graham, N.A. et al. Glucose deprivation activates a metabolic and signaling amplification loop leading to cell death. Mol Syst Biol 8, 589 (2012).

10. Sies, H. et al. Defining roles of specific reactive oxygen species (ROS) in cell biology and physiology. Nat Rev Mol Cell Biol 23, 499–515 (2022).

11. de Almeida, A. et al. ROS: Basic Concepts, Sources, Cellular Signaling, and its Implications in Aging Pathways. Oxid Med Cell Longev 2022, 1225578 (2022).

12. Qiu, X., Brown, K., Hirschey, M.D., Verdin, E. & Chen, D. Calorie restriction reduces oxidative stress by SIRT3-mediated SOD2 activation. Cell Metab 12, 662–667 (2010).

13. Cipriano, A. et al. Mechanisms, pathways and strategies for rejuvenation through epigenetic reprogramming. Nat Aging 4, 14–26 (2024).

14. Yang, J.H. et al. Loss of epigenetic information as a cause of mammalian aging. Cell 186, 305–326 e327 (2023).

15. Li, T.Y. et al. The transcriptional coactivator CBP/p300 is an evolutionarily conserved node that promotes longevity in response to mitochondrial stress. Nat Aging 1, 165–178 (2021).

16. Labbadia, J. & Morimoto, R.I. Repression of the Heat Shock Response Is a Programmed Event at the Onset of Reproduction. Mol Cell 59, 639–650 (2015).

17. Goldstein, I. et al. Transcription factor assisted loading and enhancer dynamics dictate the hepatic fasting response. Genome Res 27, 427–439 (2017).

18. Lopez-Otin, C., Blasco, M.A., Partridge, L., Serrano, M. & Kroemer, G. Hallmarks of aging: An expanding universe. Cell 186, 243–278 (2023).

19. Timmons, L. & Fire, A. Specific interference by ingested dsRNA. Nature 395, 854 (1998).

20. Kamath, R.S. & Ahringer, J. Genome-wide RNAi screening in Caenorhabditis elegans. Methods 30, 313–321 (2003).

21. Van Gilst, M.R., Hadjivassiliou, H. & Yamamoto, K.R. A Caenorhabditis elegans nutrient response system partially dependent on nuclear receptor NHR-49. Proc Natl Acad Sci U S A 102, 13496–13501 (2005).

22. Pryor, R. et al. Host-Microbe-Drug-Nutrient Screen Identifies Bacterial Effectors of Metformin Therapy. Cell 178, 1299–1312 e1229 (2019).

23. Bennett, C.F. et al. Transaldolase inhibition impairs mitochondrial respiration and induces a starvation-like longevity response in Caenorhabditis elegans. PLoS Genet 13, e1006695 (2017).

24. Li, Y., Ding, W., Li, C.Y. & Liu, Y. HLH-11 modulates lipid metabolism in response to nutrient availability. Nat Commun 11, 5959 (2020).

25. Ramachandran, P.V. et al. Lysosomal Signaling Promotes Longevity by Adjusting Mitochondrial Activity. Dev Cell 48, 685–696 e685 (2019).

26. Yuzyuk, T., Fakhouri, T.H., Kiefer, J. & Mango, S.E. The polycomb complex protein mes-2/E(z) promotes the transition from developmental plasticity to differentiation in C. elegans embryos. Dev Cell 16, 699–710 (2009).

27. Mutlu, B. et al. Distinct functions and temporal regulation of methylated histone H3 during early embryogenesis. Development 146 (2019).

28. Agger, K. et al. UTX and JMJD3 are histone H3K27 demethylases involved in HOX gene regulation and development. Nature 449, 731–734 (2007).

29. Fisher, K., Southall, S.M., Wilson, J.R. & Poulin, G.B. Methylation and demethylation activities of a C. elegans MLL-like complex attenuate RAS signalling. Dev Biol 341, 142–153 (2010).

30. Shi, Y. & Mello, C. A CBP/p300 homolog specifies multiple differentiation pathways in Caenorhabditis elegans. Genes Dev 12, 943–955 (1998).

31. Wiles, E.T. & Selker, E.U. H3K27 methylation: a promiscuous repressive chromatin mark. Curr Opin Genet Dev 43, 31–37 (2017).

32. Pasini, D. et al. Characterization of an antagonistic switch between histone H3 lysine 27 methylation and acetylation in the transcriptional regulation of Polycomb group target genes. Nucleic Acids Res 38, 4958–4969 (2010).

33. Jin, C. et al. Histone demethylase UTX-1 regulates C. elegans life span by targeting the insulin/IGF-1 signaling pathway. Cell Metab 14, 161–172 (2011).

34. Settembre, C. et al. TFEB controls cellular lipid metabolism through a starvation-induced autoregulatory loop. Nat Cell Biol 15, 647–658 (2013).

35. Hanzelmann, S., Castelo, R. & Guinney, J. GSVA: gene set variation analysis for microarray and RNA-seq data. BMC Bioinformatics 14, 7 (2013).

36. Creyghton, M.P. et al. Histone H3K27ac separates active from poised enhancers and predicts developmental state. Proc Natl Acad Sci U S A 107, 21931–21936 (2010).

37. Yu, G., Wang, L.G., Han, Y. & He, Q.Y. clusterProfiler: an R package for comparing biological themes among gene clusters. OMICS 16, 284–287 (2012).

38. Wu, T. et al. clusterProfiler 4.0: A universal enrichment tool for interpreting omics data. Innovation (Camb) 2, 100141 (2021).

39. Huang da, W., Sherman, B.T. & Lempicki, R.A. Systematic and integrative analysis of large gene lists using DAVID bioinformatics resources. Nat Protoc 4, 44–57 (2009).

40. Sherman, B.T. et al. DAVID: a web server for functional enrichment analysis and functional annotation of gene lists (2021 update). *Nucleic Acids Res* **50**, W216-W221 (2022).

41. Cheung, E.C. & Vousden, K.H. The role of ROS in tumour development and progression. Nat Rev Cancer 22, 280–297 (2022).

42. Hansen, M. et al. A role for autophagy in the extension of lifespan by dietary restriction in C. elegans. PLoS Genet 4, e24 (2008).

43. Eruslanov, E. & Kusmartsev, S. Identification of ROS using oxidized DCFDA and flow-cytometry. Methods Mol Biol 594, 57–72 (2010).

44. Lee, S.J., Hwang, A.B. & Kenyon, C. Inhibition of respiration extends C. elegans life span via reactive oxygen species that increase HIF-1 activity. Curr Biol 20, 2131–2136 (2010).

45. Castello, P.R., Drechsel, D.A. & Patel, M. Mitochondria are a major source of paraquat-induced reactive oxygen species production in the brain. J Biol Chem 282, 14186–14193 (2007).

46. Rosca, M.G. et al. Oxidation of fatty acids is the source of increased mitochondrial reactive oxygen species production in kidney cortical tubules in early diabetes. Diabetes 61, 2074–2083 (2012).

47. Song, S. et al. Molecular basis for antioxidant enzymes in mediating copper detoxification in the nematode Caenorhabditis elegans. PLoS One 9, e107685 (2014).

48. Senchuk, M.M. et al. Activation of DAF-16/FOXO by reactive oxygen species contributes to longevity in long-lived mitochondrial mutants in Caenorhabditis elegans. PLoS Genet 14, e1007268 (2018).

49. Lee, H. et al. The Caenorhabditis elegans AMP-activated protein kinase AAK-2 is phosphorylated by LKB1 and is required for resistance to oxidative stress and for normal motility and foraging behavior. J Biol Chem 283, 14988–14993 (2008).

50. Millan-Zambrano, G., Burton, A., Bannister, A.J. & Schneider, R. Histone post-translational modifications - cause and consequence of genome function. Nat Rev Genet 23, 563–580 (2022).

51. Balsalobre, A. & Drouin, J. Pioneer factors as master regulators of the epigenome and cell fate. Nat Rev Mol Cell Biol 23, 449–464 (2022).

52. Zhong, M. et al. Genome-wide identification of binding sites defines distinct functions for Caenorhabditis elegans PHA-4/FOXA in development and environmental response. PLoS Genet 6, e1000848 (2010).

53. Hu, Q. et al. BLMP-1 is a critical temporal regulator of dietary-restriction-induced response in Caenorhabditis elegans. Cell Rep 43, 113959 (2024).

54. Mango, S.E., Lambie, E.J. & Kimble, J. The pha-4 gene is required to generate the pharyngeal primordium of Caenorhabditis elegans. Development 120, 3019–3031 (1994).

55. Weiss, E.P. et al. Lower extremity muscle size and strength and aerobic capacity decrease with caloric restriction but not with exercise-induced weight loss. J Appl Physiol (1985) 102, 634–640 (2007).

56. Lakowski, B. & Hekimi, S. The genetics of caloric restriction in Caenorhabditis elegans. Proc Natl Acad Sci U S A 95, 13091–13096 (1998).

57. Wu, L. et al. An Ancient, Unified Mechanism for Metformin Growth Inhibition in C. elegans and Cancer. Cell 167, 1705–1718 e1713 (2016).

58. Onken, B. & Driscoll, M. Metformin induces a dietary restriction-like state and the oxidative stress response to extend C. elegans Healthspan via AMPK, LKB1, and SKN-1. PLoS One 5, e8758 (2010).

59. Honjoh, S., Yamamoto, T., Uno, M. & Nishida, E. Signalling through RHEB-1 mediates intermittent fasting-induced longevity in C. elegans. Nature 457, 726–730 (2009).

60. Galloy, M. et al. Ubiquitination of the histone variant mH2A1.2 prevents toxic RAD18 accumulation at a subset of genomic loci upon replication stress. Mol Cell (2025).

61. Brenner, S. The genetics of Caenorhabditis elegans. Genetics 77, 71–94 (1974).

62. Concordet, J.P. & Haeussler, M. CRISPOR: intuitive guide selection for CRISPR/Cas9 genome editing experiments and screens. Nucleic Acids Res 46, W242–W245 (2018).

63. Dickinson, D.J., Pani, A.M., Heppert, J.K., Higgins, C.D. & Goldstein, B. Streamlined Genome Engineering with a Self-Excising Drug Selection Cassette. Genetics 200, 1035–1049 (2015).

64. Escorcia, W., Ruter, D.L., Nhan, J. & Curran, S.P. Quantification of Lipid Abundance and Evaluation of Lipid Distribution in Caenorhabditis elegans by Nile Red and Oil Red O Staining. J Vis Exp (2018).

65. Mukhopadhyay, A., Deplancke, B., Walhout, A.J. & Tissenbaum, H.A. Chromatin immunoprecipitation (ChIP) coupled to detection by quantitative real-time PCR to study transcription factor binding to DNA in Caenorhabditis elegans. Nat Protoc 3, 698–709 (2008).

66. Andrews, S. (2010).

67. Ewels, P., Magnusson, M., Lundin, S. & Kaller, M. MultiQC: summarize analysis results for multiple tools and samples in a single report. Bioinformatics 32, 3047–3048 (2016).

68. Bushnell, B., Rood, J. & Singer, E. BBMerge - Accurate paired shotgun read merging via overlap. PLoS One 12, e0185056 (2017).

69. Patro, R., Duggal, G., Love, M.I., Irizarry, R.A. & Kingsford, C. Salmon provides fast and bias-aware quantification of transcript expression. Nat Methods 14, 417–419 (2017).

70. Martin, F.J. et al. Ensembl 2023. *Nucleic Acids Res* **51**, D933–D941 (2023).

71. Huber, W. et al. Orchestrating high-throughput genomic analysis with Bioconductor. Nat Methods 12, 115–121 (2015).

72. Soneson, C., Love, M.I. & Robinson, M.D. Differential analyses for RNA-seq: transcript-level estimates improve gene-level inferences. F1000Res 4, 1521 (2015).

73. Love, M.I., Huber, W. & Anders, S. Moderated estimation of fold change and dispersion for RNA-seq data with DESeq2. Genome Biol 15, 550 (2014).

74. Benjamini, Y. & Hochberg, Y. Controlling the False Discovery Rate - a Practical and Powerful Approach to Multiple Testing. J Roy Stat Soc B Met 57, 289–300 (1995).

75. Walker, M.D. et al. WormPaths: Caenorhabditis elegans metabolic pathway annotation and visualization. Genetics 219 (2021).

76. Sarasija, S. & Norman, K.R. Measurement of ROS in Caenorhabditis elegans Using a Reduced Form of Fluorescein. Bio Protoc 8 (2018).

77. Zhou, Y. et al. A secreted microRNA disrupts autophagy in distinct tissues of Caenorhabditis elegans upon ageing. Nature Communications 10, 4827 (2019).

78. Cedillo, L. et al. Ether lipid biosynthesis promotes lifespan extension and enables diverse pro-longevity paradigms in Caenorhabditis elegans. Elife 12 (2023).

