## Supplemental Figures for "Hormetic nutrient stress promotes longevity by orchestrating histone acetylation on key lipid catabolism and antioxidant defense genes"

### Supplementary Information

### Supplementary Information contains seven figures.

Extended Data Fig. 1 Zhou et al.

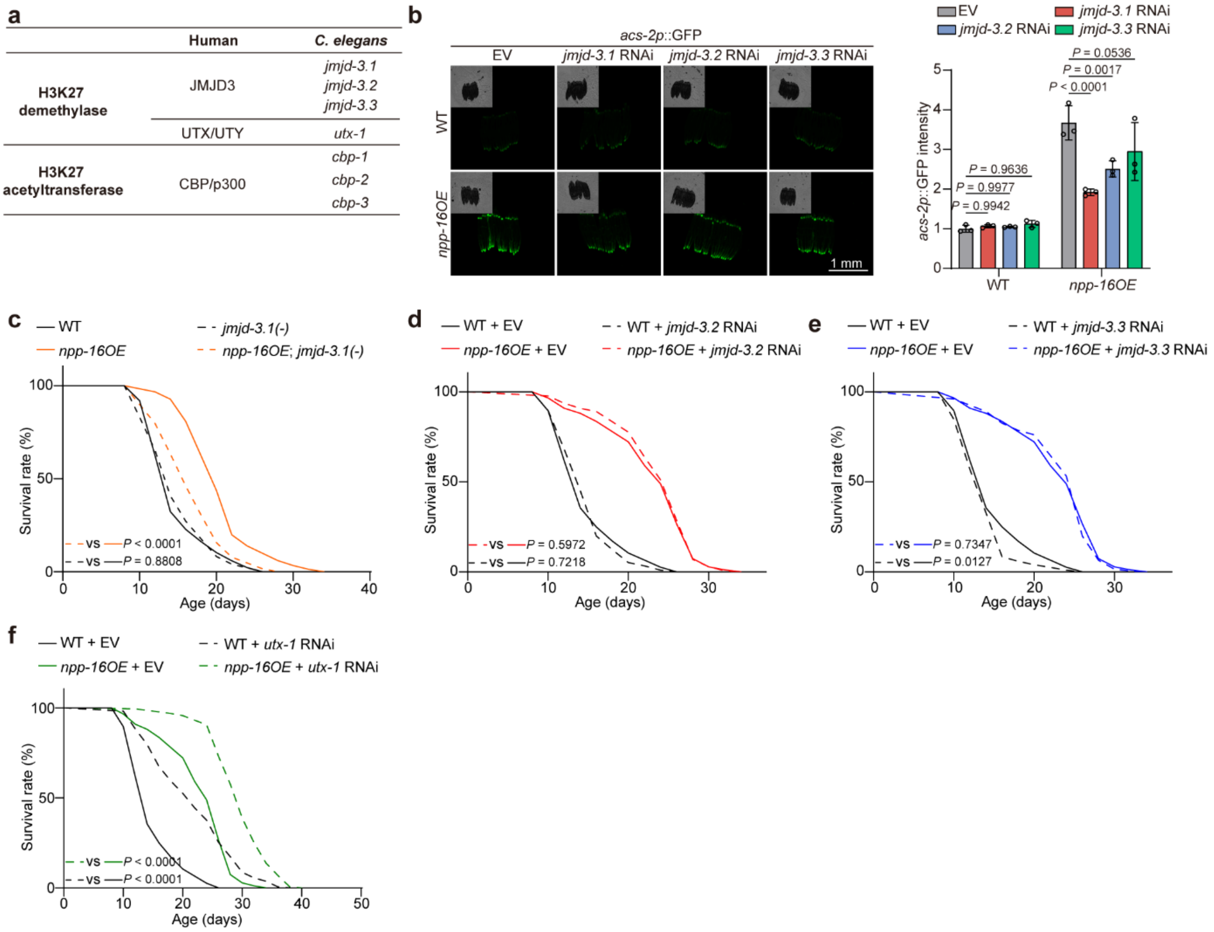


**Extended Data Fig. 1 JMJD-3.1 is the main H3K27 demethylase responsible for lifespan extension in worms overexpressing NPP-16. a**, Orthologues of H3K27 demethylases JMJD3 and UTX/UTY, and acetyltransferase CBP/p300 in *C. elegans*. **b**, The activated *acs-2p::GFP* reporter in *npp-16OE* worms is suppressed by RNAi knockdown of JMJD3 family members, in which the effect of *jmjd-3.1* RNAi is more robust than the other two members. Scale bar: 1 mm. n=3 independent experiments. **c**, *jmjd-3.1(-)* mutation partially rescues the lifespan extension by activating NPP-16. **d**,**e**, *jmjd-3.2* and *jmjd-3.3* RNAi knockdown does not affect the lifespan extension in *npp-16OE* worms. **f**, The longevity-promoting effect of *utx-1* RNAi knockdown is parallel to the lifespan extension in *npp-16OE* worms. See also Supplementary Data 3 for independent biological replicates and summary lifespan statistics. Error bars represent SD. Statistical significance was calculated by two-way ANOVA (**b**) and Mantel-Cox Log Rank (**c**-**f**).

Extended Data Fig. 2 Zhou et al.

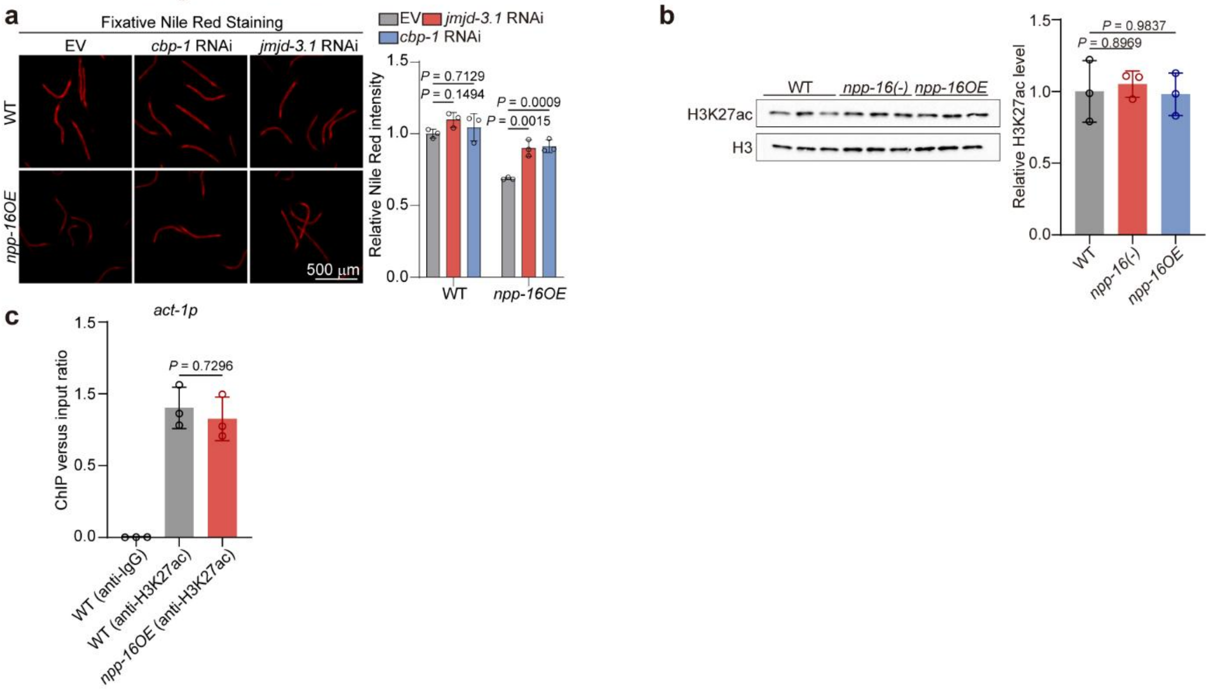


**Extended Data Fig. 2 Knockdown of individual H3K27ac modulators increases the lipid storage by NPP-16 activation, whereas NPP-16 has less impact on the global H3K27ac level. a**, *jmjd-3.1*, and *cbp-1* RNAi knockdown partially rescue the fat mass reduction in *npp-16OE worms*. n=3 independent experiments. Scale bar: 500 μm. **b**, *npp-16(-)* mutation and *npp-16OE* barely impact the global level of H3K27ac in worms. H3 serves as an internal control. n=3 independent experiments. **c**, ChIP-qPCR analyses reveal that H3K27ac is unchanged by *npp-16OE* at the promoter of *act-1* (*act-1p*). n=3 independent experiments. Error bars represent SD. Statistical significance was calculated by two-way ANOVA (**a**) and one-way ANOVA (**b**,**c**).

Extended Data Fig. 3 Zhou et al.

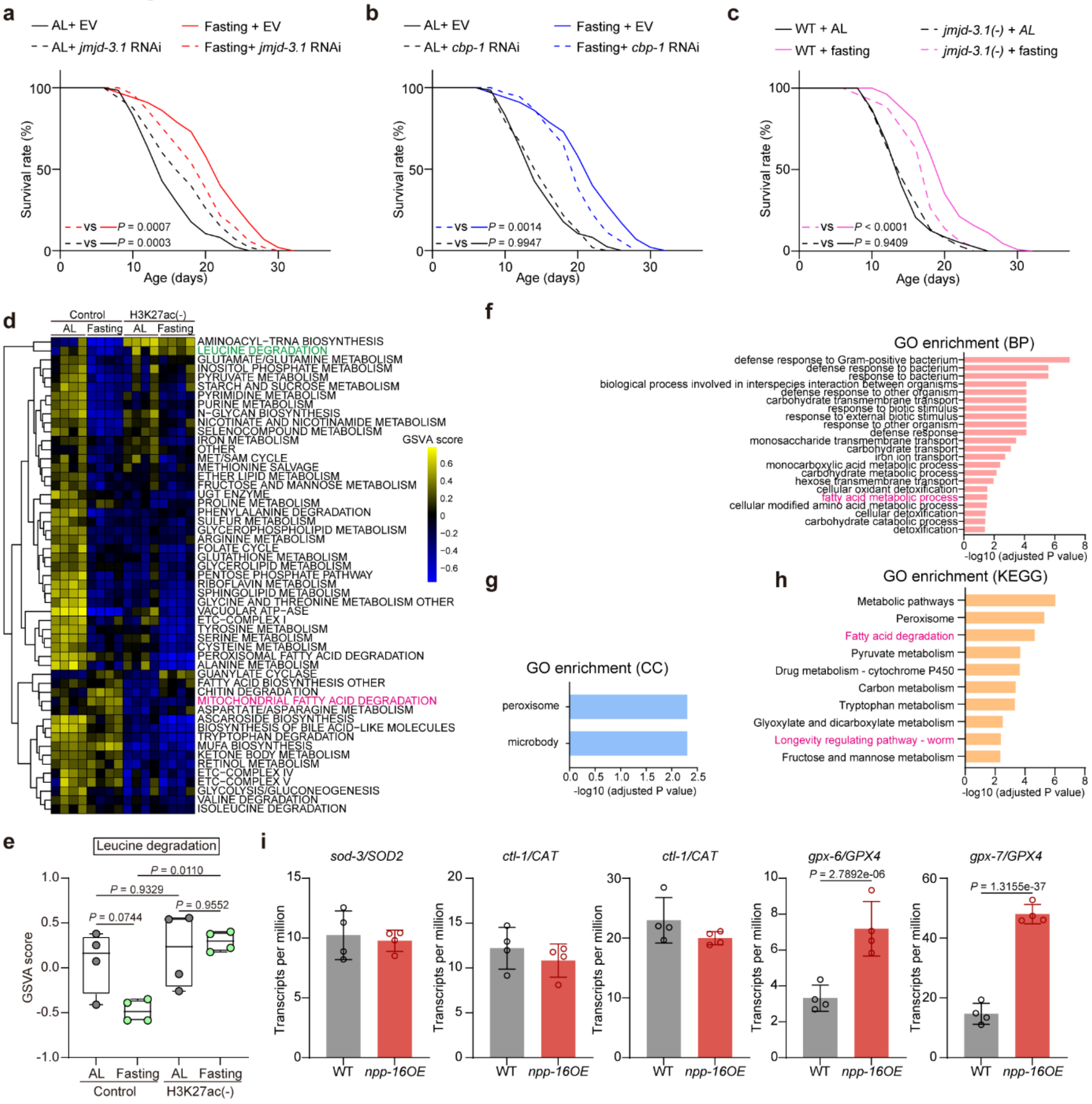


**Extended Data Fig. 3 H3K27ac modulators are crucial for the transcriptomic alterations and longevity in response to nutrient stress. a-b**, *jmjd-3.1* and *cbp-1* RNAi knockdown partially rescues the lifespan extension of hormetic fasting. **c**, *jmjd-3.1(-)* mutation suppresses the lifespan extension of hormetic fasting partially. **d-e**, The gene set variation analysis (GSVA) of RNA-seq data, showing that leucine degradation and mitochondrial fatty acid degradation are inhibited and activated by fasting in a H3K27ac-dependent manner, respectively. **f-h**, GO-term enrichment of DEGs up-regulated by fasting and depending on H3K27ac, showing that processes and pathways relevant to lipid catabolism and longevity are activated by fasting in a H3K27ac-dependent manner. BP, biological process. CC, cellular component. KEGG, Kyoto encyclopedia of genes and genomes. **i**, *gpx-6* and *gpx-7* are significantly increased by *npp-16OE* in the previous RNA-seq dataset. See Supplementary Data 2 for detailed statistics from RNA-seq (**d-h**). See also Supplementary Data 3 for independent biological replicates and summary lifespan statistics. Error bars represent SD. Statistical significance was calculated by Mantel-Cox Log Rank (**a-c**), empirical Bayes testing with Benjamini-Hochberg correction for multiple comparisons using limma() (**e**), and DESeq2 Wald tests with Benjamini-Hochberg FDR correction (**i**).

**Extended Data Fig. 4 Zhou et al.**

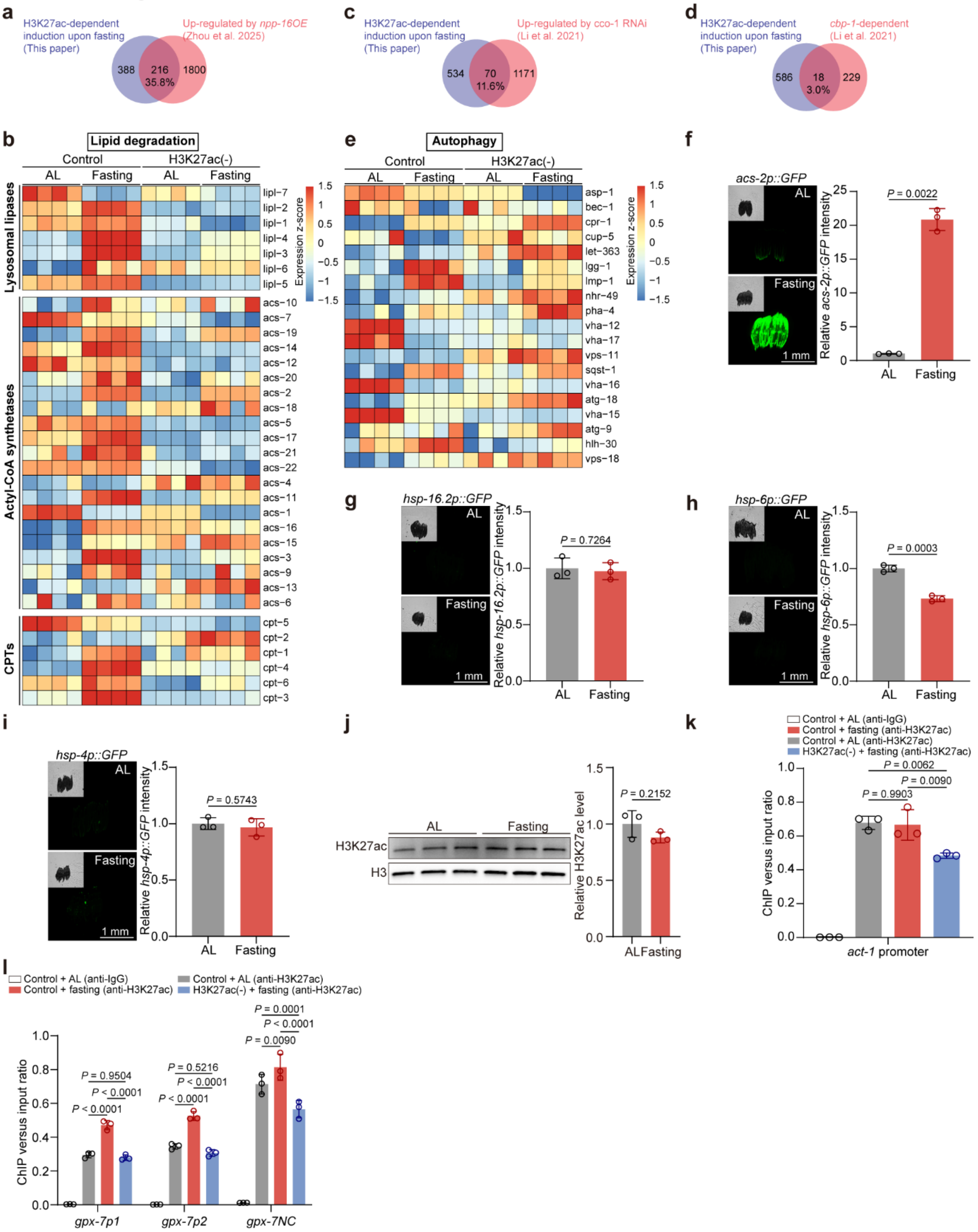


**Extended Data Fig. 4 Hormetic nutrient stress activates lipid catabolism specifically without altering other stress responses. a**, A Venn diagram showing the number of differentially expressed genes (DEGs) upregulated by fasting depending on H3K27ac, and the overlapping DEGs increased by *npp-16OE* as previously reported. **b**, A heat map indicates the expression of lysosomal lipases, Actyl-CoA synthetases, and carnitine palmitoyl transferase (CPTs) in RNA-seq data. **c,d**, Venn diagrams showing the number of DEGs comparing this study with published RNA-seq datasets. **e**, A Heat map showing the expression of indicated genes associated with autophagy in RNA-seq data. **f**, *acs-2p::GFP* reporter is significantly induced by fasting for 24 hours at D1. n=3 independent experiments. **h-i**, *hsp-16.2p::GFP*, *hsp-6p::GFP*, and *hsp-4p::GFP* reporters are unchanged by fasting for 24 hours at D1. Scale bars: 1 mm. n=3 independent experiments. **j**, The global H3K27ac level is unchanged by hormetic fasting at D1. H3 serves as an internal control. n=3 independent experiments. **k**, H3K27ac level at *act-1* promoter is unchanged by fasting but inhibited in H3K27ac(-) worms. n=3 independent experiments. **l**, ChIP-qPCR analyses reveal that the elevated H3K27ac signal at *gpx-7* promoter by fasting is blunted in H3K27ac(-) worms. The flanking sequences at or near the 3'-UTR serve as the negative controls (*gpx-7NC*) for ChIP-qPCR. n=3 independent experiments. Error bars represent SD. Statistical significance was calculated by unpaired *t*-test (**f-j**), one-way ANOVA (**k**), and two-way ANOVA (**l**).

Extended Data Fig. 5 Zhou et al.

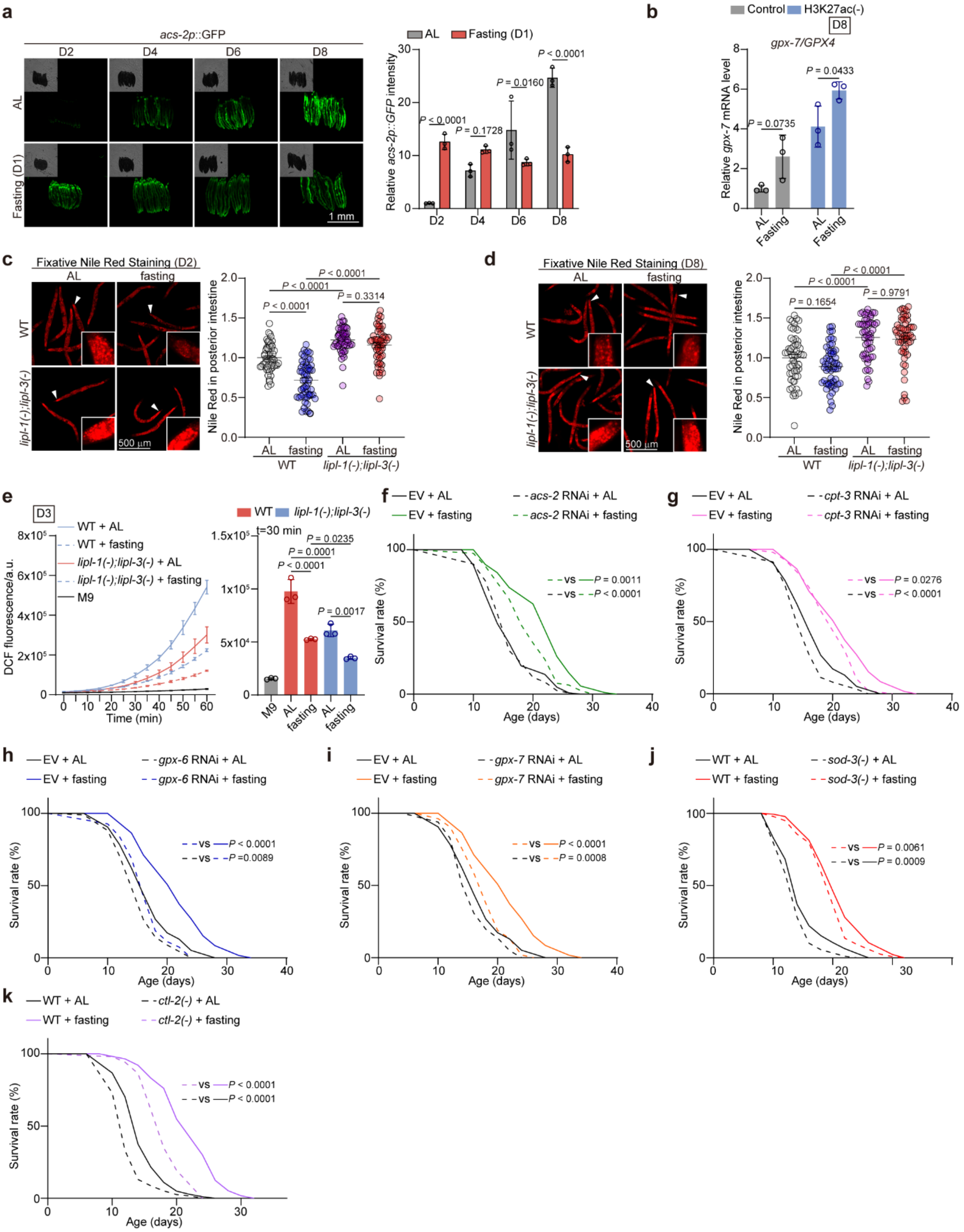


**Extended Data Fig. 5 Inhibition of lysosomal lipases suppresses fat mass reduction and ROS generation in response to hormetic fasting.**

**a**, *acs-2p::GFP* reporter is significantly activated by fasting for 24 hours at D1, which is eliminated by refeeding to D8. Scle bar: 1 mm. n=3 independent experiments. **b**, *gpx-*

*7/GPX4* mRNA expression at D8 is induced in worms subjected to fasting at D1, which is unchanged by H3K27ac inhibition. n=3 independent experiments. **c,** Neutral lipid storage indicated by fixative Nile red staining reveals that *lipl-1(-);lipl-3(-)* double mutation blocks the fat mass consumption after fasting for 24 hours at D1. Scale bar: 500 μm. n=3 independent experiments. **d**, The reduced fat mass resulting from fasting at D1 is recovered to the level compared to AL feeding at D8. Scale bar: 500 μm. n=3 independent experiments. **e**, *lipl-1(-);lipl-3(-)* mutation reduces the ROS level in worms subjected to AL and fasting. n=3 independent experiments. n=3 independent experiments. **f-k**, The lifespan of worms subject to hormetic fasting at D1 for 24 hours in indicated mutants or treated with indicated RNAi. See also Supplementary Data 2 for independent biological replicates and summary lifespan statistics. Error bars represent SD (**a**, **b**, and **e**) and SEM (**c** and **d**). Statistical significance was calculated by two-way ANOVA (**a-b**), one-way ANOVA (**c-e**), and Mantel-Cox Log Rank (**f-k**).

Extended Data Fig. 6 Zhou et al.

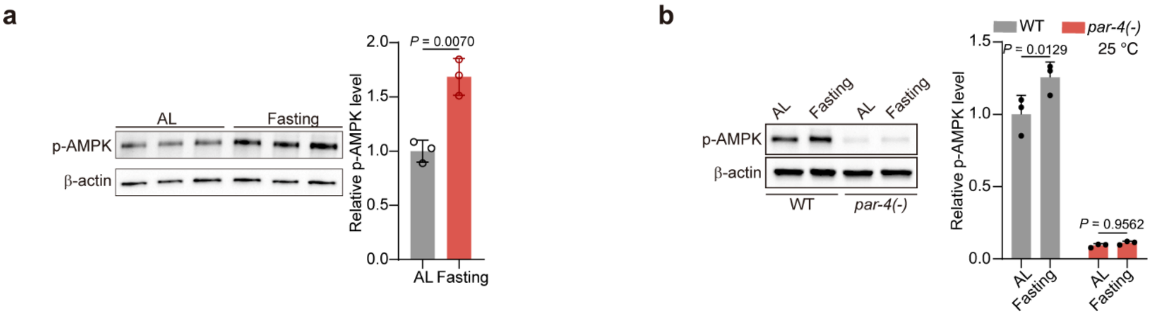


**Extended Data Fig. 6 PAR-4/LKB1 is required for AMPK activation in response to fasting.**

**a**, Phosphorylation level of AMPK (p-AMPK) is elevated by hormetic fasting for 24 hours at D1. n=3 independent experiments. **b**, *par-4(-)* mutation ablates the induced p-AMPK in response to hormetic fasting. β-actin serves as an internal control. n=3 independent experiments. Error bars represent SD. Statistical significance was calculated by unpaired *t*-test (**a**) and two-way ANOVA (**b**).

Extended Data Fig. 7 Zhou et al.

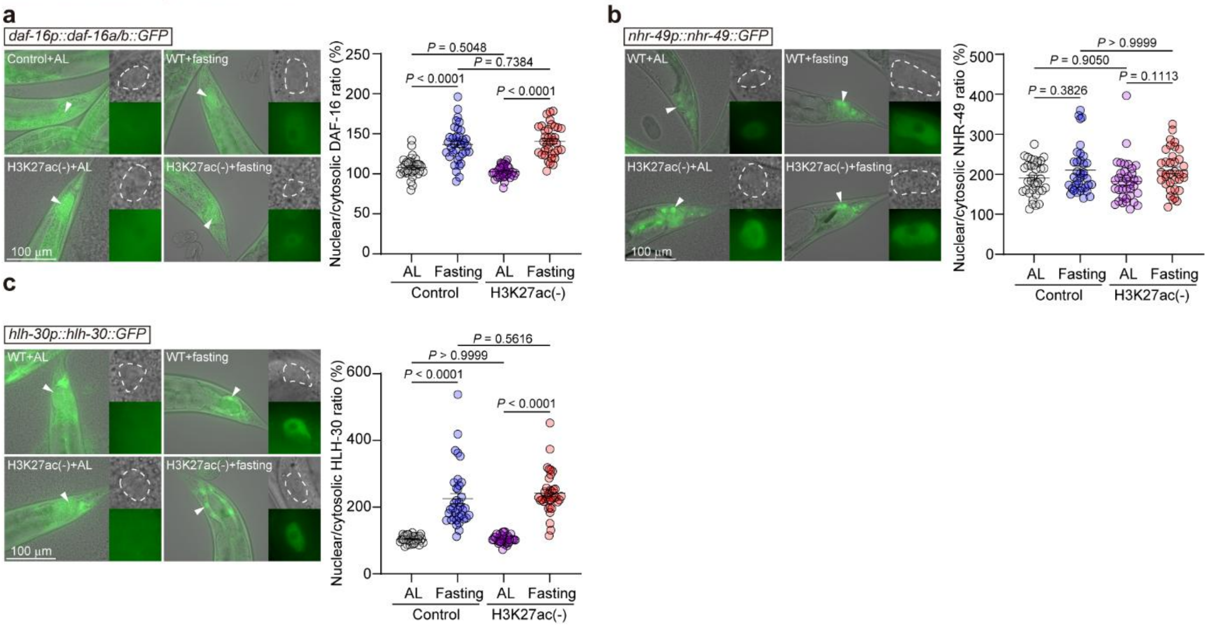


**Extended Data Fig. 7 Nuclear localization of DAF-16, NHR-49, and HLH-30 is unchanged when H3K27ac is inhibited.**

**a-c**, Representative images and quantification of overexpressed DAF-16 (**a**), NHR-49(**b**), and HLH-30 (**c**) in WT and H3K27ac(-) worms in response to fasting for 6 hours at D1. n=3 independent experiments, containing 36 worms in total. Error bars represent SEM. Scale bar: 100 μm. Statistical significance was calculated by one-way ANOVA.
